# Dual Noncanonical Amino Acid Incorporation Enabling Chemoselective Protein Modification at Two Distinct Sites in Yeast

**DOI:** 10.1101/2022.10.19.512873

**Authors:** Priyanka Lahiri, Meghan S. Martin, Briana R. Lino, Rebecca A. Scheck, James A. Van Deventer

## Abstract

Incorporation of more than one non-canonical amino acid (ncAA) within a single protein endows the resulting construct with multiple useful features such as augmented molecular recognition or covalent crosslinking capabilities. Herein, for the first time, we demonstrate the incorporation of two chemically distinct ncAAs into proteins biosynthesized in *Saccharomyces cerevisiae*. To complement ncAA incorporation in response to the amber (TAG) stop codon in yeast, we evaluated opal (TGA) stop codon suppression using three distinct orthogonal translation systems. We observed selective TGA readthrough without detectable cross-reactivity from host translation components. Readthrough efficiency at TGA was modulated by factors including the local nucleotide environment, gene deletions related to the translation process, and the identity of the suppressor tRNA. These observations facilitated systematic investigation of dual ncAA incorporation in both intracellular and yeast-displayed protein constructs, where we observed efficiencies up to 6% of wildtype protein controls. The successful display of doubly-substituted proteins enabled the exploration of two critical applications on the yeast surface - A) antigen-binding functionality; and B) chemoselective modification with two distinct chemical probes through sequential application of two bioorthogonal click chemistry reactions. Lastly, by utilizing a soluble form of a doubly-substituted construct, we validated the dual incorporation system using mass spectrometry and demonstrated the feasibility conducting selective labeling of the two ncAAs sequentially using a ”single-pot” approach. Overall, our work facilitates the addition of a 22^nd^ amino acid to the genetic code of yeast and expands the scope of applications of ncAAs for basic biological research and drug discovery.

**Graphical Abstract:** 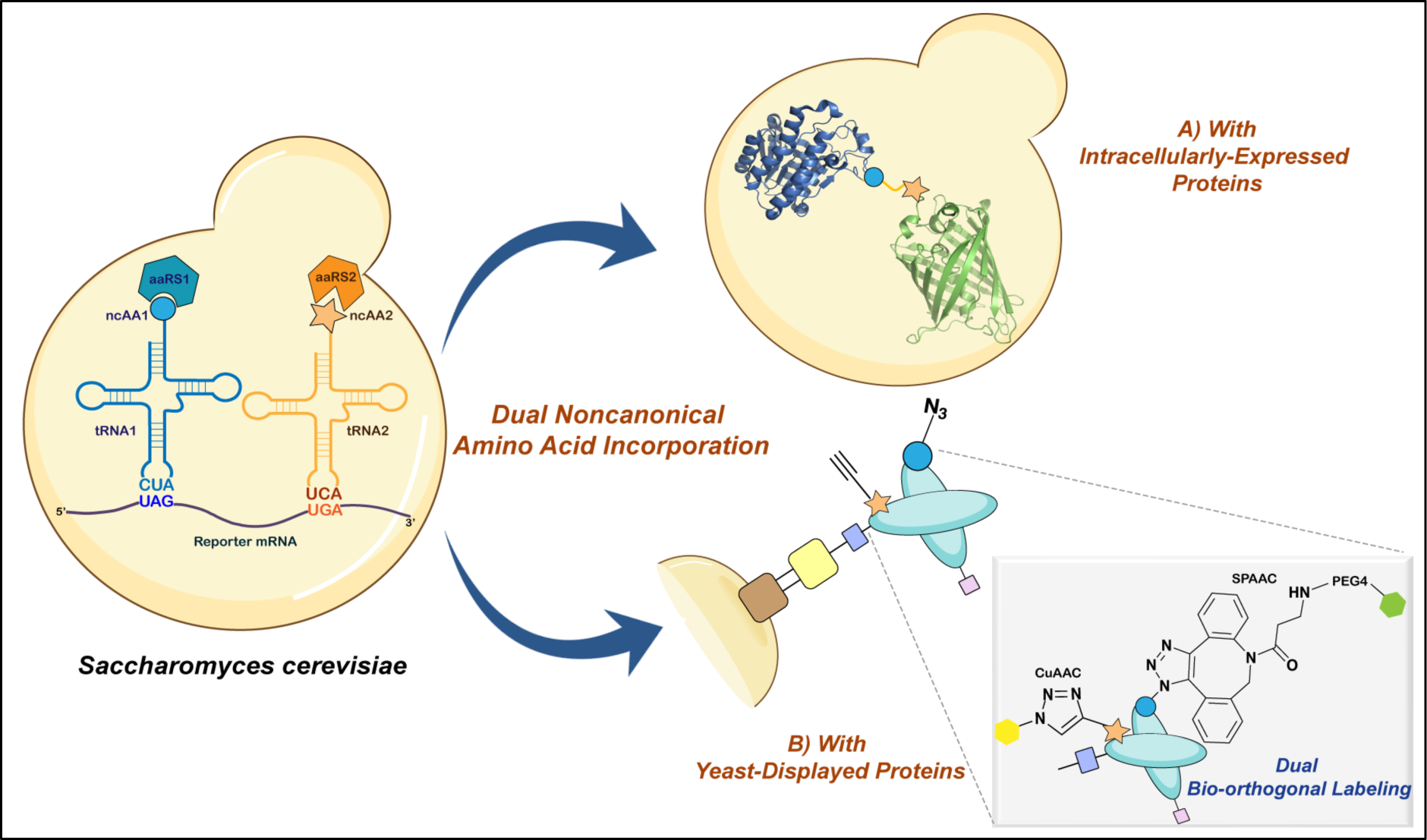

Herein we report the establishment of dual noncanonical amino acid incorporation in yeast to support expression and site-selective labelling of doubly substituted proteins in solution and on the yeast surface.

## Introduction

The genetic encoding of noncanonical amino acids (ncAAs) in proteins allows for precise manipulation of their native structures and the introduction of new properties and chemical entities^1–7^. The number of ncAAs that can be encoded during protein biosynthesis and the organisms capable of encoding them is constantly expanding^8, 9^. The most successful dual ncAAs incorporation systems have been demonstrated in bacterial and mammalian expression systems; these approaches have enabled a wide range of applications such as exploring protein dynamics, protein-protein crosslinking, and mimicking epigenetic modifications^10–18^. However, to the best of our knowledge, such dual ncAA incorporation systems have not yet been implemented in *S. cerevisiae* or any other yeast. With reasonably fast growth kinetics and biological machinery analogous to mammalian cells*, S. cerevisiae* is a critical model organism for understanding fundamentals of eukaryotic biology, functions of disease-implicated genes as well as serving as workhorse for generating complex recombinant proteins^19–21^. Additionally, powerful genetic, genomic, and engineering tools are now available in yeast, including single-gene knockout collections, strains with synthetic chromosomes, and yeast display systems^22–26^. Therefore, engineering a dual ncAA incorporation system in *S. cerevisiae* would enable applications in areas of protein engineering, chemical biology, synthetic biology, and other related disciplines^27^.

Incorporating two distinct ncAAs in cells requires the use of two orthogonal codons and two corresponding aminoacyl-tRNA-synthetase/tRNA pairs (aaRS/tRNA pairs); also referred to as orthogonal translation systems (OTSs). These pairs must be orthogonal to one another as well as to the host cell’s native translation machinery. Such orthogonal codons can be repurposed sense codons, nonsense (“stop”) codons, or engineered frameshift codons. For both bacteria and mammalian cells, these orthogonal codons have been discovered and utilized in various combinations, but in yeast, only the amber (TAG) codon has been repurposed for ncAA incorporation^11–13, 16, 17, 28–31^. The lack of additional validated orthogonal codons and mutually-orthogonal translation systems in yeast has hindered the development of dual ncAA incorporation systems, in contrast to systems in bacteria and mammalian cells that support the decoding of multiple orthogonal codons with structurally and functionally diverse _ncAAs_11,16,30,32–35.

Efforts are underway in yeast to expand the range of available OTSs for genetic code manipulation. Recent studies by He *et al* and Tan *et al* demonstrated the isolation of *E. coli* aaRS variants that support translation with new ncAAs including sulfotyrosine^36, 37^. Parallel efforts from our lab have also led to the discovery of a broad range of *E. coli* tyrosyl-tRNA synthetase (*Ec*TyrRS) and *E. coli* Leucyl-tRNA synthetase (*Ec*LeuRS) variants with a variety of ncAA incorporation properties, including new-to-yeast ncAAs and improved selectivity for known ncAAs^38^. Similar efforts by Chatterjee and coworkers in engineered *E. coli* strains has led to the identification of additional *Ec*aaRS variants derived from *Ec*TyRS and *Ec*TrpRS with unique substrate preferences for use in mammalian cells^33, 39^. While not yet evaluated in yeast, such *Ec*aaRS variants have the potential to further broaden the range of OTSs available for use in this organism. Beyond *E. coli*-derived OTSs, translation machinery derived from archaeal Pyrrolysyl-tRNA synthetases (PylRS)-tRNA pairs continues to mature^10, 40–45^. PylRS-based OTSs from several organisms and their engineered derivatives support diverse ncAA incorporation and efficient decoding of a range of orthogonal codons. We and others have advanced the use of such PylRS-tRNA pairs in yeast, which presents opportunities for the identification of mutually orthogonal translation systems^46–49^. The growing versatility of OTSs available in yeast sets the stage for pursuing dual ncAA incorporation in this important model organism.

Herein, for the first time, we report the systematic establishment of dual ncAA incorporation systems in *S. cerevisiae* suitable for applications involving both intracellular and yeast-displayed proteins. We repurpose the opal (TGA) nonsense codon to function alongside the amber (TAG) nonsense codon in yeast and demonstrate the successful decoding of these two orthogonal codons within the same mRNA transcript by different combinations of mutually orthogonal OTSs. Our detailed investigations identify potential non-cognate interactions that can interfere with the fidelity of the system and favorable genetic conditions augmenting the decoding efficiencies of both TAG and TGA codons. This engineered dual ncAA incorporation system enables site-specific incorporation of distinct ncAAs pairs, allowing for the generation of doubly-labeled proteins through sequential strain-promoted azide-alkyne cycloaddition (SPAAC) and copper-catalyzed azide-alkyne cycloaddition (CuAAC) reactions on the yeast surface. These reactions and corroborating investigations on soluble proteins demonstrate the incorporation and selective addressability of two distinct chemical functionalities within proteins biosynthesized in yeast.

## Results

### A second orthogonal codon identified in yeast

For complementing the TAG codon as a location for ncAA incorporation in yeast, another nonsense codon—opal (TGA) codon—was selected due to its low usage within yeast genome and prior use for ncAA incorporation in mammalian systems^17, 50–53^. However, for TGA to be repurposed as an orthogonal codon in yeast, it must fulfill two important criteria: *i*) Orthogonality; indicating that TGA should not be decoded at high levels by the endogenous translational machinery or non-cognate OTSs; and *ii*) Decoding efficiency; meaning that TGA should be efficiently decoded only by OTSs that utilize tRNAUCAs and not by OTSs utilizing tRNACUAs or other exogenous machinery. As there are examples of natural TGA readthrough in yeast by endogenous translation machineries under selected conditions (which also holds true for TAG), it is critical that the second criteria is met to minimize aberrant readthrough^54^. Thus, to systematically evaluate both criteria at TGA codons, a dual-plasmid approach was used, where the OTS was encoded on one plasmid and the reporter protein bearing the orthogonal codon was encoded on another. This dual-plasmid approach allows for the parallel assessment of different cognate and non-cognate interactions: aaRS:tRNA, aaRS:ncAA, aaRS:cAA (canonical amino acid), and codon:anticodon (Figure 1A).

**Figure 1.**
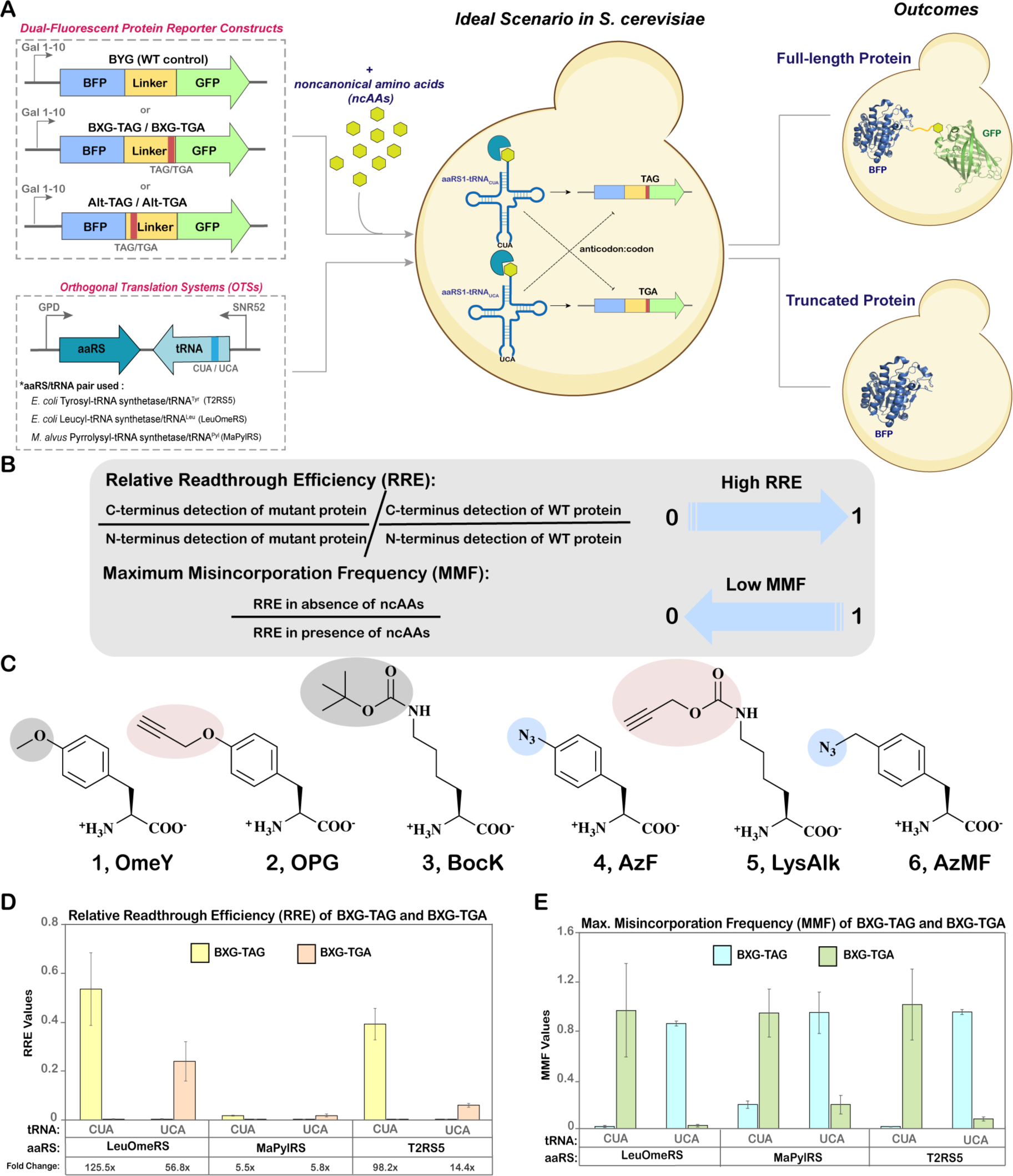
Evaluation of TAG versus TGA stop codon suppression. A) Schematic overview of dual plasmid approach outlining the architecture of dual-fluorescent reporter constructs and orthogonal translation system (OTS) constructs. The possible cognate and non-cognate codon:anticodon interactions of amber and opal suppressor tRNAs are shown as bold arrows and dotted lines, respectively. B) Equations used for determining relative readthrough efficiency (RRE) and maximum misincorporation frequency (MMF), quantitative parameters for evaluating ncAA incorporation efficiency and fidelity, respectively. C) Structures of ncAAs used in this study. Shaded regions highlight functional groups, some of which can undergo specific bioorthogonal reactions. **1**, *O*-methyl-L-tyrosine (OmeY); **2**, *N*^ε^-Boc-L-Lysine (BocK); **3**, *p*-propargyloxyl-L-phenylalanine (OPG); **4**, *p*-azido-L-phenylalanine (AzF); **5**, *N*^ε^-propargyloxycarbonyl-L-lysine (LysAlk); **6**, 4-azidomethyl-L-phenylalanine (AzMF). D) Relative Readthrough Efficiency (RRE) of BXG-TAG and BXG-TGA reporters with indicated tRNA and aaRS constructs calculated from flow cytometry experiments. The fold changes reported below the plots (e.g. 125.5x) indicate the fold change in readthrough between BXG-TAG and BXG-TGA by a particular OTS; these values were calculated from the ratios of median fluorescence intensities (MFIs) of readthrough events of these reporter constructs (the larger value was used as the numerator). E) Maximum Misincorporation Frequency (MMF) calculated for BXG-TAG and BXG-TGA reporter constructs. Flow cytometry dot plots and and bar graphs of MFIs for all data in this figure are depicted in Figure S1 and Figure S2, respectively. The error bars in D and E were derived from measurements of biological triplicates, starting with standard deviations of MFI values and propagated using error propagation equations (during calculations of RRE and MMF).

Employing our established, galactose-inducible dual fluorescent protein reporter system, we evaluated the incorporation of ncAAs at TGA and TAG codons^55, 56^. The reporter system is a fusion protein containing a N-terminal fluorescent protein (BFP) and a C-terminal fluorescent protein (GFP) connected by a linker with desired orthogonal codon at a permissive site. The orthogonal codon was either a TAG (referred to as BXG-TAG) or TGA (referred to as BXG-TGA); a “wild-type” linker containing no stop codon served as a positive control (referred to as BYG; Figure 1A and Table S4)^68^. This dual-fluorescent reporter system enabled the use of the quantitative metrics Relative Readthrough Efficiency (RRE) and Maximum Misincorporation Frequency (MMF) to evaluate characteristics of protein translation with ncAAs (Figure 1B). RRE quantifies the readthrough of an orthogonal codon with respect to wildtype protein synthesis, with values typically ranging between 0 and 1. An RRE value of 0 indicates that all proteins produced are truncated at the orthogonal codon, while an RRE value of 1 indicates readthrough efficiency equivalent to wildtype protein synthesis. MMF measures the probability of unintentional canonical amino acid incorporation at the orthogonal codon, with values approaching zero during high fidelity ncAA incorporation and one or higher as the misincorporation frequency increases. An ideal scenario representing efficient and site-selective ncAA incorporation should have a high RRE and a low MMF value.

We utilized variants of three yeast-compatible OTSs: *E. coli* tyrosyl tRNA synthetase/tRNA^Tyr^, *E. coli* leucyl tRNA synthetase/tRNA^Leu^ and *M. alvus* pyrrolysyl tRNA synthetase/tRNA^Pyl^, and cloned each of them separately into the pRS315 plasmid backbone under constitutive promoters (Figure 1A). Recognition of TAG or TGA codons during mRNA translation was facilitated by mutating the anticodon sequences of suppressor tRNAs to CUA or UCA, respectively. This simple approach facilitated evaluation of cognate and non-cognate aminoacylation events to determine whether the aaRSs can tolerate changes in anticodons, as anticodon identity is known to modulate tRNA recognition and aminoacylation in many aaRSs^57^. For the *Ec*LeuRS/tRNA^Leu^ pair, we used the polyspecific aaRS variant LeuOmeRS with a T252A mutation in its editing domain and employed the ncAA *O*-methyl-L-tyrosine (**1**; OmeY) as a substrate^58^. For the *Ec*TyrRS/tRNA^Tyr^ pair, we employed a polyspecific aaRS variant, T2RS5, identified through our prior high-throughput screening and used it with the ncAA *p*-propargyloxyl-L-phenylalanine (**2**; OPG) as a substrate^38^. Lastly, for the PylRS/tRNA^Pyl^ pair, we used the *M. alvus* PylRS with the ncAA *N^ε^-*Boc-L-Lysine (**3**; BocK) as a substrate (Figure 1C)^61^.

For our initial investigation of ncAA incorporation at TGA codons, we conducted a series of flow cytometry experiments to determine RRE and MMF at “cognate” and “noncognate” orthogonal codons for each of the OTSs (Figure 1D). The three aaRS/tRNA pairs encoding plasmids with either tRNACUA or tRNAUCA were separately co-transformed into the *S. cerevisiae* strain RJY100 with a reporter construct encoding a cognate or a non-cognate orthogonal codon, or a WT reporter (absence of orthogonal codons; Figure 1A)^59^. All transformants were induced either in the absence or in the presence of 1 mM cognate ncAA and subjected to flow cytometry analysis and RRE and MMF calculations (Figure 1A; Figure S1). Notably, all three OTSs supported ncAA incorporation in response to TGA when the OTSs contained tRNAs with complementary UCA anticodon, but not a noncognate anticodon. Additionally, quantitative analyses indicated high RRE and low MMF values for all cognate OTS:reporter combinations tested, and the opposite trend for non-cognate OTS:reporter combinations. These data demonstrated that these OTSs exhibit specificity at the codon:anticodon interaction level, without any detectable cross-reactivity (Figure 1C and 1D). Moreover, both flow cytometry dot plots and median fluorescence intensity (MFI) graphs revealed insignificant background fluorescence levels for both TAG and TGA codons in the absence of any ncAAs, suggesting negligible participation of endogenous translation machinery in codon readthrough events (Figure S1 and S2). On direct comparison with TAG, TGA was decoded 2-6 fold less efficiently by both T2RS5/tRNAUCA^Tyr^ and LeuOmeRS/tRNAUCA^Leu^ pairs (Figure 1C). The observation of lower levels of TGA readthrough by the two *E. coli* OTSs in comparison to TAG readthrough is consistent with the prior observations of Chatterjee and coworkers with TGA and TAG readthrough in mammalian cells^17^. In contrast, for the *Ma*PylRS/tRNA^Pyl^ pair, the decoding efficiencies of both TAG and TGA codons were at equivalent levels when paired with tRNAs containing the “cognate” anticodon. However, RRE values determined with these systems were observed to be indistinguishable from samples lacking ncAAs. Hence, to further substantiate the translational activities of these samples, we evaluated median fluorescence intensities of C-terminal reporter expression and observed clear differences in fluorescence between samples induced in the absence and presence of ncAAs (Figure S2). Overall, these data validated TGA as an orthogonal codon in yeast for site-specific ncAA incorporation, with no detectable cross-reactivity with TAG at the codon:anticodon level.

### Sequence context impacts TGA suppression efficiency

Prior to combining constructs encoding both TAG and TGA codons for dual ncAA incorporation, we sought to better understand the underlying factors that can influence ncAA incorporation in response to TGA. Previous studies have shown that nucleotides upstream and downstream of TAG codons can affect ncAA incorporation efficiency in bacteria, mammalian, and yeast expression systems^28, 56, 60–63^. These observations and the fact that natural TGA readthrough events in yeast (with a cAA) are also dictated by their neighboring nucleotides, motivated us to explore whether the same trend holds true for ncAA incorporation at TGA codons^50, 64^. For our evaluation, we selected an alternate location within the linker of our reporter construct, referred to as ‘Alt-TAG’, which in our prior work demonstrated moderate but significantly decreased readthrough across different OTS and ncAA combinations^56^. We replaced the TAG in this construct with TGA to generate a new construct, referred to as Alt-TGA (Figure 1A, Table S4). For quantifying solely the changes related to orthogonal codon position, other experimental parameters including OTS identities, induction conditions, and ncAA types were all kept identical to the experimental parameters used in combination with BXG reporters.

Flow-cytometric analysis revealed similar ncAA incorporation efficiency and fidelity trends at the Alt-TGA position as at the BXG-TGA position, irrespective of the OTS used (Figure 2A, S3, and S4A). For both *E. coli* OTSs, ncAA incorporation in response to TAG was more efficient than incorporation in response to TGA, while for the *Ma*PylRS/tRNA^Pyl^ OTS, ncAA incorporation in response to either TAG or TGA (with corresponding “cognate” tRNAs) was comparable; this is evident in evaluations of RRE, MMF, and median fluorescence intensities of C-terminal GFP fluorescence levels (Figure 2A and S4). Interestingly, comparing RRE values between the two stop codon positions (BXG-*versus* Alt-reporters) confirmed that the overall readthrough at the position within the Alt-construct was substantially lower than the corresponding readthrough at the position within the BXG-construct. This corroborated our and other research groups’ findings that the nucleotide environment around the orthogonal codon impacts the readthrough efficiency in yeast, and suggests that this trend holds irrespective of whether the readthrough occurs at a TAG or TGA codon^56^. To further assess changes in readthrough performance that occur between the two stop codon positions, we determined the ratios of the MFIs of GFP fluorescence for cognate OTS:reporter combinations at the BXG or BXG-TGA position to the Alt-TAG or Alt-TGA position (Figure 2B). Surprisingly, when comparing changes in readthrough at these two positions, the resulting fold changes in TGA readthrough across the three OTSs was distinct in comparison to corresponding fold changes in TAG. Theoretically, the fold change for a particular nonsense codon should not change between OTSs if the nucleotide context of the reporter is the sole contributing factor to readthrough efficiency. We attribute the observed differences in TGA readthrough (in comparison to TAG readthrough) to the anticodon change in the tRNA and to differences in the nucleotide sequences (and presumably identity elements) of the tRNAs in each OTS. Therefore, our data indicates that efficient codon:anticodon interactions are governed by specific features of both the stop codon sequence context in the reporter and the nucleotide sequence of the tRNA component of the OTS.

**Figure 2.**
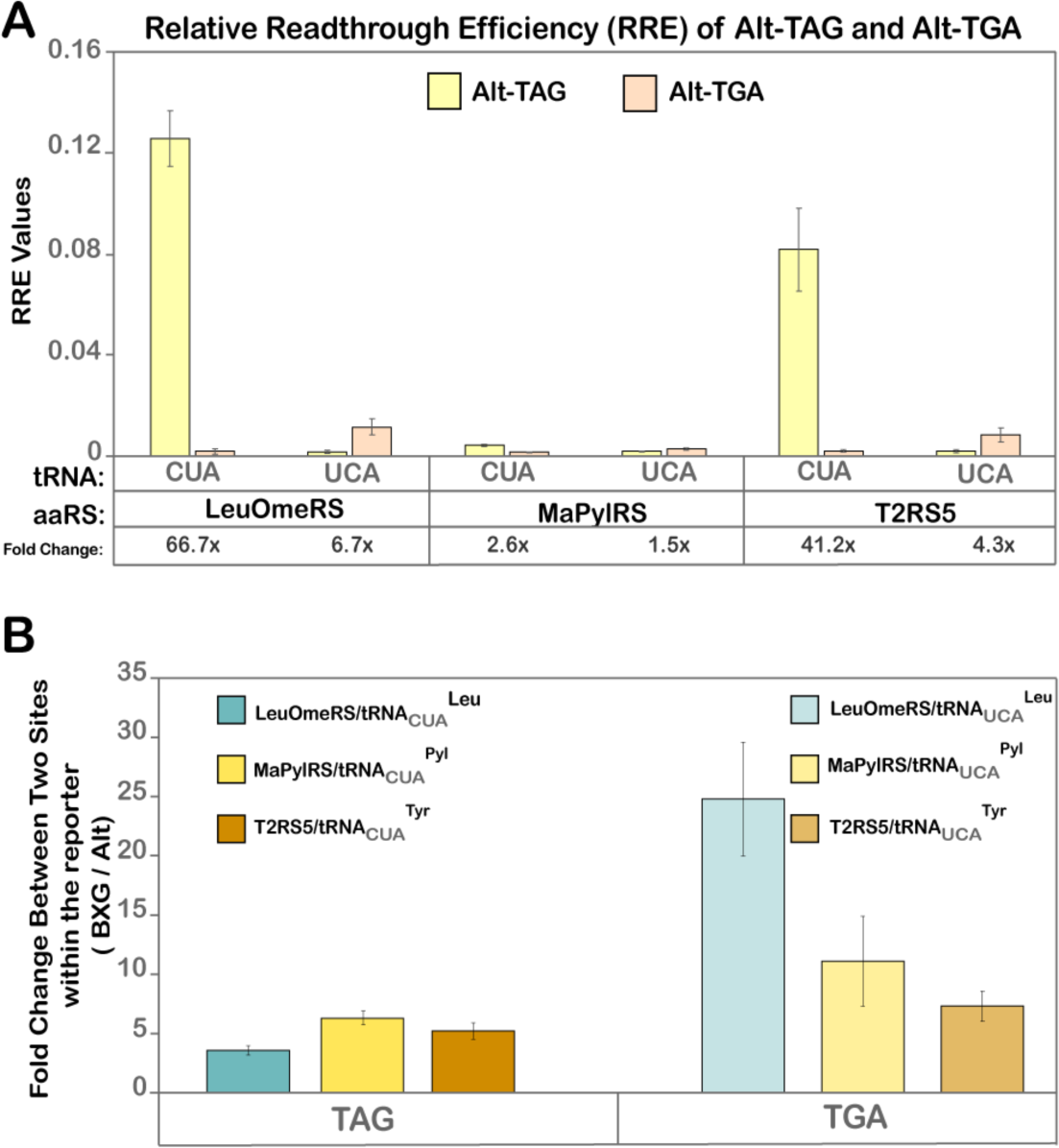
Quantitative analysis of readthrough events for Alt-TAG and Alt-TGA constructs. A) Relative Readthrough Efficiency (RRE) for Alt-TAG and Alt-TGA reporter constructs calculated from MFI values obtained from flow cytometry dot plot data, depicted in Figure S3. The fold-changes in readthrough (eg: 66.7X) in the table below indicate the fold change in readthrough between Alt-TAG and Alt-TGA by a particular OTS; these values were calculated from the ratios of MFIs of readthrough events of these reporter constructs (the larger value was used as the numerator). The corresponding MMF graph and the MFI bar graph are represented in Figure S4. B) Fold change in readthrough performance between the two codon positions in the reporter (BXG and Alt) obtained from the ratio of MFI values of cognate OTS:reporter interaction at the BXG-TAG (or BXG-TGA) site to the Alt-TAG (or Alt-TGA) site. The error bars in both A and B were calculated from the standard deviation of MFI values and then propagated in subsequent calculations using error propagation equations.

### Single-gene knockout yeast strains enhance TGA readthrough efficiency

Numerous engineering strategies are known to augment stop codon readthrough, including single-gene deletions, whole-genome synthesis, and engineering various components of the translation machinery^1, 26, 30, 38, 65, 66^. Prior work from our group showed that single-gene knockout strains of *S. cerevisiae* improved ncAA incorporation efficiencies in response to the TAG codon for various OTSs and ncAAs combinations ^56, 66, 67^. Based on our past work, we narrowed investigations here to two single-gene knockout yeast strains, *ppq1τι* and *tpa1τι*; deletions of these genes are also known to enhance nonsense suppression with near-cognate tRNAs^68^. *TPA1* regulates mRNA poly(A) tail length and interacts with eukaryotic release factors eRF1 and eRF3, while *PPQ1* is a serine/threonine phosphatase with an undefined role in translation termination^69, 70^. We employed a BY4741 yeast strain background due to the ready availability of single-gene knockouts in this background. This necessitated transferring the BXG- and Alt-reporter constructs from a pCTCON2-plasmid backbone (*TRP1* marker) to a pRS416-plasmid backbone (*URA3* marker), since *trp1* is not an available auxotrophic marker in BY4741. Otherwise, all experimental conditions were kept identical to the conditions used above, and we included TAG-containing reporter constructs for side-by-side comparison of readthrough effects in *ppq1τι* and *tpa1τι* strains.

NcAA incorporation efficiencies were evaluated via flow cytometry for all strains using each of the 3 OTSs and 2 stop codon positions summarized in Figure 1A. Our analysis (both RRE and MFIs) revealed that the *ppq1τι* strain demonstrated statistically significant (2-3 fold) enhancements in TGA readthrough across different OTSs and codon positions compared to the WT BY4741 strain (Figure 3, Figure S9). These improvements were more readily apparent for the Alt-TGA (and Alt-TAG) constructs than for the corresponding BXG constructs (Figure 3; Figure S5-S7, S9). On other hand, for the *tpa1Δ* strain, slight increases in readthrough efficiency were only observed in some cases where *Ec*TyrRS/tRNA^Tyr^ or *Ec*LeuRS/tRNA^Leu^ OTSs were used. In the case of *Ma*PylRS/tRNA^Pyl^, no enhancements in TGA readthrough efficiencies were observed in *tpa1Δ* under the conditions tested. Under all conditions tested here, improvements in ncAA incorporation efficiency in knockout strains were not accompanied by detectable changes to the apparent fidelity of ncAA incorporation, as evidenced by the lack of changes in MMF (Figure S8-S9). The overall trends in RRE values determined in BY4741 strains were consistent with the trends observed in RJY100. Most notably, RRE values for ncAA incorporation at TGA codons were lower than RRE values at TAG codons under corresponding conditions (Figure 1D and 3A; Figure 2A and 3B). Interestingly, when changing the plasmid backbone from pCTCON2 to pRS416, there was a modest decrease in RRE values with LeuOmeRS/tRNA^Leu^ OTSs for both TAG and TGA, but not for the other two OTSs. These observations are consistent with our previous findings, indicating that changes in plasmid backbone can impact the apparent performance of some OTSs for reasons that have not yet been elucidated^56^. In summary, the *ppq1Δ* strain enhanced ncAA incorporation across both orthogonal codons, independent of the codon position or OTS type.

**Figure 3.**
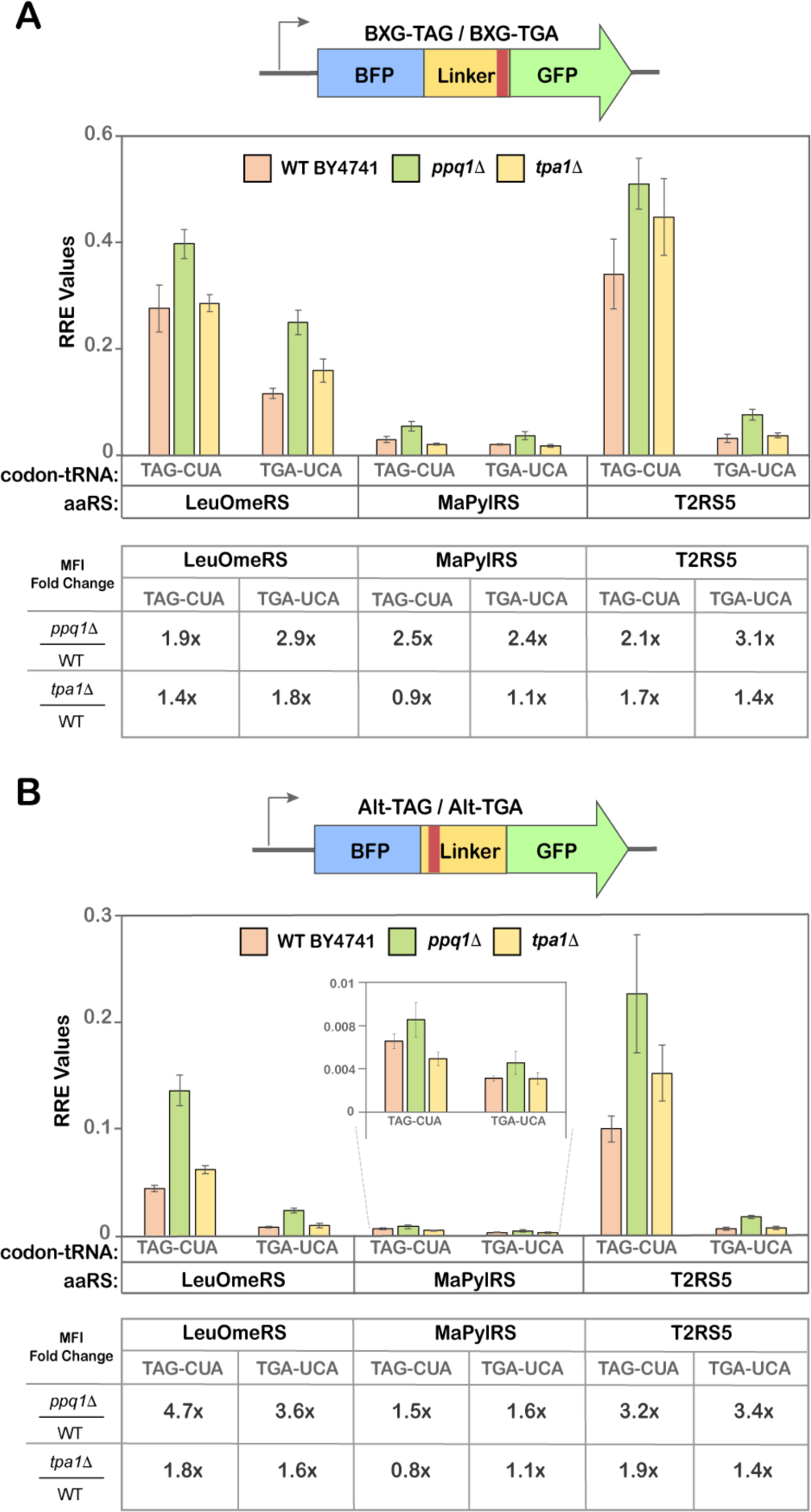
Quantitative readthrough measurements in single-gene knockout yeast strains. A) Relative Readthrough Efficiency (RRE) of BXG-TAG and BXG-TGA reporters with indicated tRNA and aaRS constructs calculated from flow cytometry experiments for the yeast strain BY4741 and single-gene knockouts *ppq1Δ* and *tpa1Δ*. B) Relative Readthrough Efficiency (RRE) of Alt-TAG and Alt-TGA reporters with indicated tRNA and aaRS constructs calculated from flow cytometry experiments for the yeast strain BY4741 and single-gene knockouts *ppq1Δ* and *tpa1Δ*. *O*-methyl-L-tyrosine (**1**; OmeY) was used as substrate for LeuOmeRS/tRNA^Leu^, *N^ε^-*Boc-L-Lysine (**2**; BocK) was used as a substrate for MaPylRS/tRNA^Pyl^; and *p*-propargyloxyl-L-phenylalanine (**3**; OPG) was used as a substrate for T2RS5/tRNA^Tyr^. The error bars are representative of three independent experiments, calculated from the standard deviations and processed through error propagation equations. The fold change in readthrough values (e.g. 1.9x) are represented in table format (below each RRE plot in A and B) and were obtained from the ratios of MFI values of a particular codon:OTS interaction in a knockout yeast strain to the corresponding MFI value in the WT BY4741 strain. The corresponding MMF graph and the MFI bar graphs for the data in this figure are depicted in supplementary Figure S8 and S9.

### Dual ncAA incorporation system for intracellular proteins

With a broad understanding of factors influencing ncAA incorporation at both TAG and TGA sites in separate constructs, we next sought to enable dual ncAA incorporation within a single protein. For facilitating expression of two OTSs, we replaced the reporter plasmid with our previously described single plasmid system encoding both a TAG-suppressing OTS and the inducible reporter system (Figure 4A)^81^. The reporter was modified to encode both TAG and TGA within the linker, at the “BXG” and “Alt” positions in both possible orientations, resulting in two mutant reporters: BX2G-AO (A: Amber (TAG); O: Opal (TGA)) and BX2G-OA. These plasmids were co-transformed with a separate plasmid encoding the second, TGA-suppressing OTS. We also used a single-plasmid version of the BYG reporter (orthogonal codons absent) as a positive control (Figure 4A). For initial analyses of dual ncAA incorporation, we selected the *Ec*TyrRS/tRNA^Tyr^ pair to suppress the TAG codon because it exhibited the highest TAG codon readthrough in the BY4741 background among all the OTSs and across different codon positions. We employed the TyrOmeRS variant in place of the T2RS5 variant because TyrOmeRS exhibits a narrower ncAA substrate preference, reducing potential crosstalk with other OTSs^71^. For suppressing TGA, we used the same OTS variants as were used in the investigations above, resulting in the following combinations of OTSs for dual ncAA incorporation experiments: *Ec*TyrRS(TyrOmeRS)/tRNACUA^Tyr^ + *Ec*LeuRS(LeuOmeRS)/tRNAUCA^Leu^ (hereafter, referred as ‘TyrRSTAG + LeuRSTGA’) and *Ec*TyrRS(TyrOmeRS)/tRNACUA^Tyr^ + *Ma*PylRS/tRNAUCA^Pyl^ (hereafter, referred as ‘TyrRSTAG+ PylRSTGA’) (Figure 4A). For these experiments, we induced BY4741 and *ppq1*τ1 yeast transformed with the dual systems under three sets of conditions: *i)* absence of ncAAs; *ii)* presence of one ncAA; and *iii)* presence of both ncAAs. These allowed systematic evaluation of cognate readthrough events as well as identification of aberrant readthrough events that may result from non-cognate interactions such as aaRSs:tRNA or aaRSs:ncAA (Figure 4A). For minimizing off-target aaRS:ncAA interactions, the ncAAs recognized most efficiently by their “cognate” OTSs were selected: 4-Azidomethyl-L-phenylalanine (**6**) as a substrate for LeuOmeRS, 4-Azido-L-phenylalanine (**4**) for TyrOmeRS and, *N^ε^*-Boc-L-Lysine (**5**) for MaPylRS.

**Figure 4.**
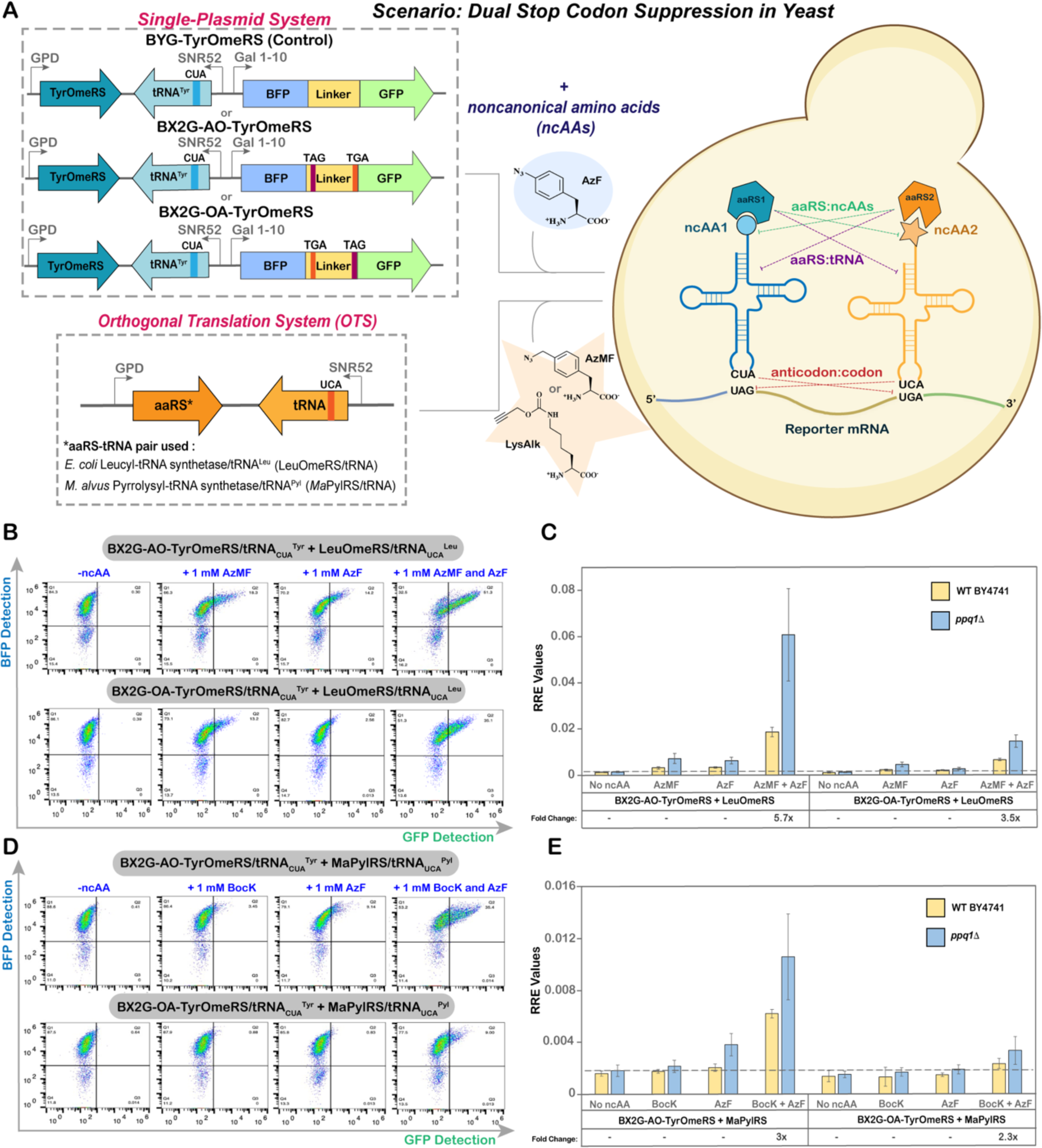
Evaluation of dual ncAA incorporation system in yeast. A) Experimental outline of dual plasmid approach to facilitate expression of two OTSs in the host cell. Dotted lines (---) and color-matched labels indicate potential non-cognate interactions that can occur. B) Flow cytometry dot plots of readthrough events of BX2G-AO and BX2G-OA in *ppq111*, employing the OTS combination of *Ec*TyrRS/tRNACUA^Tyr^ + *Ec*LeuRS/tRNAUCA^Leu^ and four different induction conditions. C) Corresponding Relative Readthrough Efficiency (RRE) calculated for OTS combination of *Ec*TyrRS/tRNACUA^Tyr^ + *Ec*LeuRS/tRNAUCA^Leu^ for the reporters BX2G-AO and BX2G-OA in BY4741 and *ppq111* strains. D) Flow cytometry dot plots of readthrough events of BX2G-AO and BX2G-OA in *ppq111*, employing the OTS combination of *Ec*TyrRS/tRNACUA^Tyr^ + *Ma*PylRS/tRNAUCA^Pyl^ and four different induction conditions. E) Corresponding Relative Readthrough Efficiency (RRE) measurements for the OTS combination of *Ec*TyrRS/tRNACUA^Tyr^ + *Ma*PylRS/tRNAUCA^Pyl^ for the reporters BX2G-AO and BX2G-OA in WT BY4741 and *ppq111* strains. The error bars are representative of three independent experiments, calculated from the standard deviation values and processed through error propagation equations. The fold change in readthrough represented in the graphs (e.g. 6x; panels C and E) are obtained from the ratios of MFI values of readthrough events in *ppq111* and BY4741 strains. Corresponding maximum misincorporation frequency (MMF) graphs can be found in Supplementary Figure S12.

Following induction under all conditions, we conducted flow cytometry to explore dual ncAA incorporation. Dot plots and RRE calculations revealed clear evidence of full-length reporter expression in the presence of both ncAAs, for both OTS combinations and both orientations of orthogonal codon positions, (BX2G-AO or BX2G-OA) (Figure 4B-E, Figure S10-S11). Quantitatively, the amount of full-length reporter detected was 1–6% of wild-type reporter expression levels (Figure 4C and 4E). As expected, readthrough efficiency in *ppq1Δ* was higher than in BY4741, improving RRE for the TyrRSTAG + LeuRSTGA combination by 3–6 fold, while the RRE for the TyrRSTAG + PylRSTGA combination was improved by 2–3 fold (Figure 4C and 4E). Readthrough levels remained at background levels in the absence of ncAAs, but low levels of full-length reporter expression (<1% of the BYG WT control reporter) were observed when induced in the presence of a single ncAA. These aberrant readthrough events were more pronounced with the TyrRSTAG + LeuRSTGA combination of OTSs, likely due to non-cognate aaRSs:tRNA or aaRSs:ncAA interactions (Figure 4B-C). This observation is in line with prior work from Zheng *et al* in mammalian cells, where it was demonstrated that mutating the *E. coli* tRNA anticodon sequence resulted in recognition by non-cognate *E. coli* aaRSs^17^. In addition, since AzF and AzMF only differ by a methylene unit, there is a possibility of either of the aaRSs recognizing the “non-cognate” but structurally related ncAA. Also, there is some evidence for low-level AzF incorporation by LeuOmeRS in our earlier work^62, 68^. In comparison to TyrRSTAG + LeuRSTGA, the TyrRSTAG + PylRSTGA OTS combination exhibited much lower RRE values for both orientations of stop codons. This is consistent with our above results and with our prior work (Figure 3)^61^. Readthrough efficiencies were also affected by the orientation of the two orthogonal codons within the reporter. Readthrough experiments with BX2G-AO led to approximately 4-fold higher levels of full-length reporter than experiments with BX2G-OA for both of the OTS combinations evaluated here. These results are in agreement with our observations of individual TAG and TGA codon readthrough events at BXG- and Alt-stop codon positions. As highlighted in Figure 2A and C, the BXG position is more permissive for orthogonal codon readthrough than the Alt position. As TGA is decoded at lower levels than TAG by the OTSs used here, having TGA at a more permissive position (BX2G-AO rather than BX2G-OA) facilitated higher overall readthrough (Figure 4C and E). In conclusion, these experiments demonstrate the feasibility of conducting dual ncAA incorporation in yeast, with the combination of TyrRSTAG + LeuRSTGA yielding up to 6% of wild-type reporter levels.

### Dual ncAA incorporation system established in yeast display format

Next, we extended our dual ncAA incorporation system to yeast-displayed proteins, as yeast-display is a powerful platform to engineer proteins and supports myriad assays for evaluating properties like affinity, stability, activity, and specificity^3, 23, 72^. We began by identifying two sites in a previously reported synthetic antibody fragment (Donkey1.1 scFv) – L93 (located in complementarity determining region L3; CDR-L3) and H54 (located in CDR-H2) for encoding ncAAs in response to TAG and TGA (Figure 5A)^3^. Both of these sites have been previously demonstrated to tolerate different ncAAs inserted in response to the TAG codon. The two codons were incorporated interchangeably at both sites, giving rise to two yeast display reporter constructs: Donkey1.1-AO and Donkey1.1-OA (Figure 5A; the nomenclature for AO and OA is the same as stated in the previous section). We employed N- and C-terminal epitope tags (HA and c-myc) for quantitative flow cytometric evaluation of protein display levels. A wild-type version of the reporter protein was used as a control and to facilitate calculation of RRE and MMF. Similar to the set of plasmids used for the intracellular dual ncAA incorporation system, a single-plasmid yeast display + ncAA incorporation system was generated by cloning a TAG-suppressing OTS upstream of the reporter in the pCTCON2 vector backbone, with TGA-suppressing OTS cloned separately into the pRS315 vector backbone. Both of the plasmids were co-transformed into yeast-display strain RJY100, which facilitates the use of induction procedures identical to those used for dual fluorescent reporters^59^. For preparation of a “mutually orthogonal” *Ec*TyrRS/tRNACUA^Tyr^ + *Ec*LeuRS/tRNAUCA^Leu^ combination, we used the TyrAcFRS variant as part of an *Ec*TyrRS/tRNACUA^Tyr^ OTS to suppress TAG, and a newly-identified variant termed LysAlkRS3 as part of the *Ec*LeuRS/tRNAUCA^Leu^ OTS to suppress TGA^38^. These OTSs were chosen based on their nonoverlapping substrate specificities, where the ncAA AzF (**4**) can be encoded with TyrAcFRS/tRNACUA^Tyr^, and the ncAA LysAlk (**5**) can be encoded with LysAlkRS3/tRNAUCA^Leu^. Importantly, choosing ncAAs with bio-orthogonal reactive handles in these experiments facilitated later investigations of double labeling strategies on the yeast surface.

**Figure 5.**
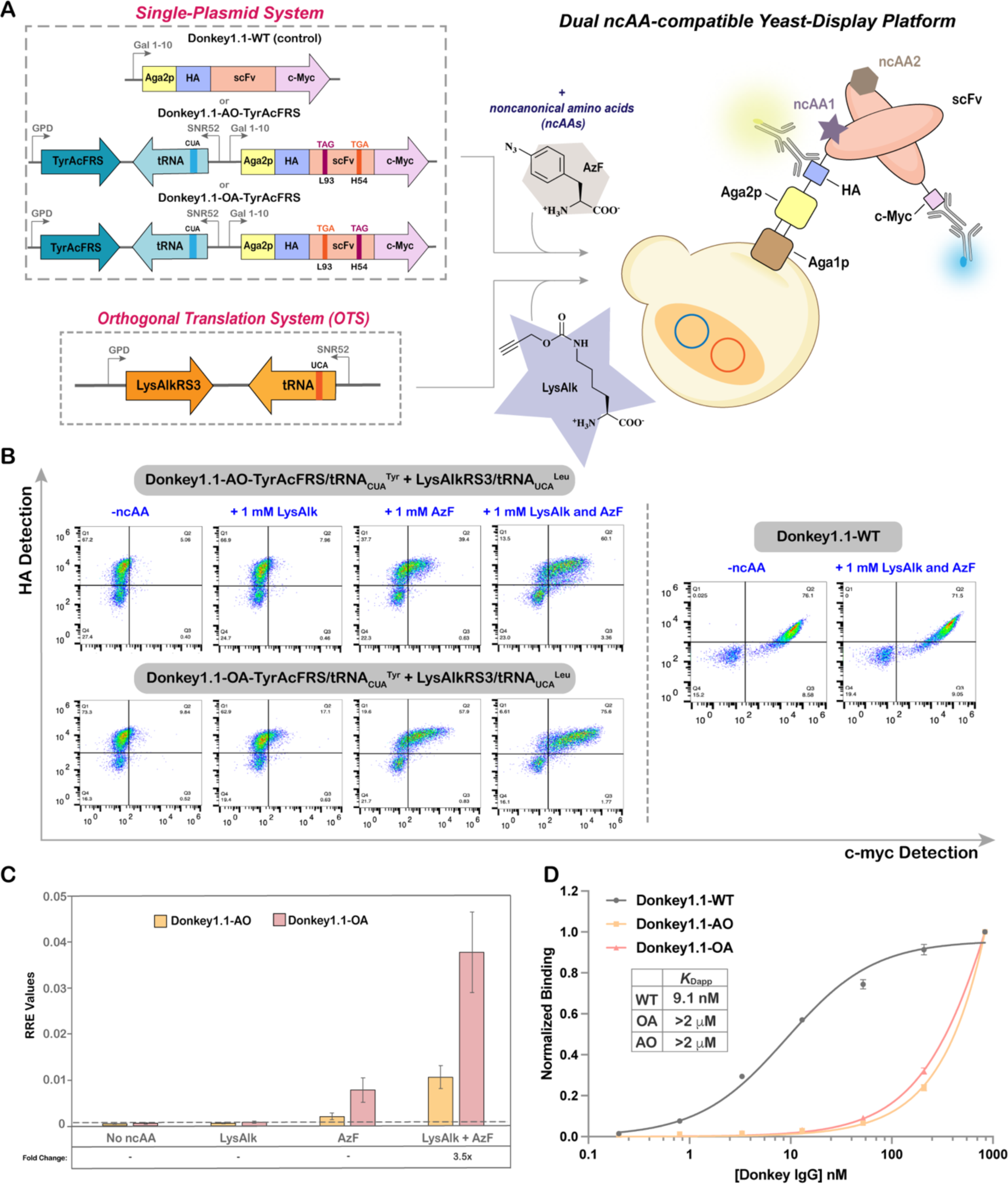
Evaluation of dual ncAA incorporation in yeast display format. A) Experimental outline for setting up a dual ncAA incorporation system with yeast-display platform. HA and c-Myc tags were employed to evaluate full-length readthrough of yeast-displayed scFvs using flow cytometry. B) Flow cytometry dot plots of readthrough experiments conducted in the RJY100 yeast display strain. Display of Donkey1.1-AO, Donkey1.1-OA and Donkey1.1 WT reporters following induction in the presence of 0, 1, or 2 ncAAs. Donkey1.1-WT was used as a positive control. C) RRE measurements comparing readthrough efficiencies between Donkey1.1-AO and Donkey1.1-OA scFv reporters across different induction conditions. Errors bars are representative of experiments conducted with three biological replicates and were determined via error propagation. The fold changes in readthrough represented in the graph (e.g. 3.5x) were obtained from the ratio of MFI values of readthrough events of Donkey1.1-OA to Donkey1.1-AO by the OTS combination of *Ec*TyrRS/tRNACUA^Tyr^ + *Ec*LeuRS/tRNAUCA^Leu^. The corresponding MMF data can be found in Supplementary Figure S13. D) Yeast display titration experiments with doubly substituted and WT synthetic antibody clones. The graph depicts normalized antigen binding levels of the three yeast-displayed scFvs obtained over a range of antigen concentrations using flow cytometry. Corresponding dot plots can be found in Supplementary Figure S14.

Flow cytometry evaluations revealed the presence of doubly-positive populations (HA+ and c-myc+) for both stop codon orientations following induction in the presence of AzF and LysAlk; this provides tangible evidence of full-length protein display on the yeast surface (Figure 5B). RRE analysis indicated that the full-length antibody display ranged between 1–4% of WT protein display levels. As observed with intracellular reporters, low levels of full-length protein (<1% of the WT protein) were also detected following induction in the presence of a single ncAA. This signal was most apparent when cultures were induced in the presence of AzF, which could be the result of non-cognate interactions between TyrAcFRS and tRNA^Leu^; LysAlkRS3 and AzF; or LysAlkRS3 and canonical amino acids^38^. A comparison of full-length protein display levels revealed that the orientation of the orthogonal codons within the Donkey1.1-OA construct leads to 3.5-fold higher full-length protein than the orientation of the orthogonal codons within the Donkey1.1-AO construct. This implies that the L93 position is more permissive for TGA readthrough than H54, consistent with our prior observation with TAG readthrough events (Figure 5C)^3^. Furthermore, we assessed the binding functionalities of the doubly-substituted yeast-displayed clones through yeast surface titration experiments and included a wild-type version of the reporter (absence of any ncAAs) as a positive control. For both doubly substituted clones, we observed clear antigen binding for concentrations as low as 3.3 nM. However, their apparent binding affinities were decreased by roughly 2 orders of magnitude in comparison to the WT protein (*K*Dapp: 9.1 nM; Figure 5D and S14). For both Donkey1.1-OA and Donkey1.1-AO, apparent *K*Ds were estimated to be in range of approximately 3–5 μM. Given that we previously observed some loss of binding affinity upon single ncAA substitution at both L93 and H54, it seems reasonable to observe further loss of binding affinity upon the introduction of a second ncAA within the same construct. Another possibility is that the low display levels of doubly substituted constructs, which result in reduced avidity on the yeast surface, decrease the apparent affinities of the constructs for the dimeric IgG antigen. Nevertheless, these data clearly demonstrate the feasibility of performing dual ncAA incorporation on the yeast surface, and further provide evidence for the retention of antigen binding functionality in doubly substituted synthetic antibody variants.

To further corroborate dual ncAA incorporation, mass spectrometry was performed with soluble forms of Donkey1.1-OA and Donkey1.1-WT. Using plasmids encoding secreted scFv-Fc constructs, we prepared and isolated pure forms of each protein via protein A affinity chromatography as determined by SDS-PAGE (Figure S15)^3^. Like yeast-displayed constructs, these secreted constructs must traverse the secretory pathway prior to secretion. Following tryptic digests, it was possible to identify the L93 and H54 peptide fragments in both WT and OA samples (Figure 6A, B). For the WT sample, the L93 fragment was observed at *m/z* of 1108.5016, *z*=4 with a neutral mass of 4429.97 (expected 4429.9849; -3.36 ppm) and the H54 fragment was observed at *m/z* of 1117.5346 *z*=2 with a neutral mass of 2233.0549 (expected 2233.0539; 0.43 ppm) (Figure 6A). For the OA sample, the L93 fragment was observed at *m/z* of 1139.2668 *z*=4 with a neutral mass of 4553.0371 (expected 4553.0533; -3.56 ppm), suggesting LysAlk incorporation, and the H54 fragment was observed at *m/z* = 1155.057 *z*=2 with a neutral mass of 2308.1015 (expected 2308.1012; 0.15 ppm) (**Figure 6B**). Thus, the observed mass change associated with ncAA incorporation in the OA sample is [M+123] for the L93 site, as expected for LysAlk incorporation. However, it is only [M+75] instead of the anticipated [M+101] for pAzF incorporation in the H54 peptide. As we and others have previously observed, pAzF is readily converted to p-aminophenylalanine (pAmF) during sample preparation and mass spectrometry analysis^38, 73^. The H54 fragment we identified is consistent (within 0.15 ppm) with the presence of pAmF, which we can use a surrogate for pAzF incorporation prior to analysis.

**Figure 6.**
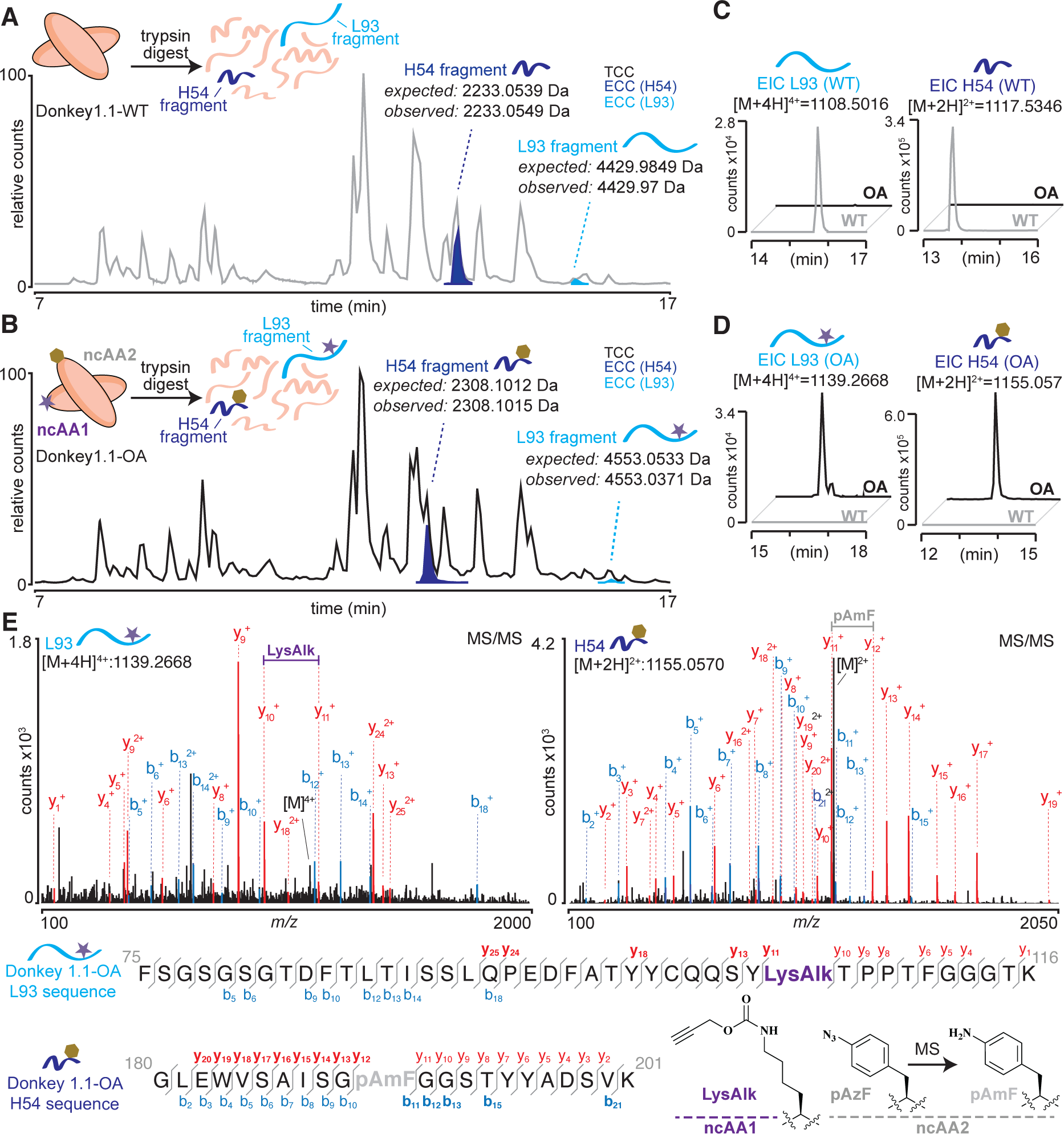
Confirmation of dual ncAA incorporation by mass spectrometry. After tryptic digestion of A) Donkey1.1-WT or B) Donkey1.1-OA, the resulting peptide fragments were analyzed by liquid-chromatography mass spectrometry (LC-MS). Total compound chromatograms (TCC) show the resulting digest mixture. Extracted compound chromatograms (ECC) show the peaks corresponding to L93 (cyan) and H54 (blue) tryptic fragments. A shift in retention time can be observed for ncAA-containing peptides in the OA sample, as compared to WT. Extracted ion chromatograms for C) WT L93 and H54 peptides or D) ncAA-containing L93 and H54 peptides show that the identified peptides are specific to either WT or OA samples. E) MS/MS analysis of ncAA-containing L93 and H54 peptides from the OA sample. Observed ions that are specific only to ncAA-containing (but not WT) peptides are shown in bold. blue, b ions; red, y ions.

Further confirmation of these assignments was accomplished by using extracted ion chromatograms (EIC) to search for specific tryptic fragments in both WT and OA samples. Using this approach, we confirmed that the unmodified (WT) H54 and L93 fragments could not be found in the OA digest (Figure 6C). Likewise, we also found that the LysAlk-containing L93 fragment and pAmF-containing H54 fragment could not be detected in the WT digest (Figure 6D). Finally, using MS/MS analysis, it was possible to identify diagnostic b+ and y+ ions confirming that both AzF (pAmF) and LysAlk were incorporated at the expected sites within the observed tryptic fragments (Figure 6E, Figures S16 and S17). Taken together, these results allow us to unambiguously confirm that Donkey1.1-OA contains both azide and alkyne functionalities imparted by dual ncAA incorporation.

### Chemoselective double labeling of yeast-displayed and soluble proteins

Covalent attachment of two distinct chemical probes to dual ncAA-substituted proteins has enabled diverse biological applications, including FRET for studying protein conformation and dynamics^12, 13, 16^. Here, we investigate, for what we believe is the first time, the combination of SPAAC and CuAAC reactions to install two different chemical probes within yeast-displayed proteins. Generally, SPAAC and CuAAC reactions are not considered mutually compatible, as they both require azide reactants. However, prior studies have demonstrated that sequential use of SPAAC and CuAAC reactions can support selective biomolecule labeling, both *in vivo* and *in vitro*^74–76^. For facilitating exploration of such sequential reactions with our system, we used the yeast-displayed dual ncAA-containing antibody reporters Donkey1.1-AO and Donkey1.1-OA (Figure 5A; see previous section) along with the corresponding wild-type control. Figure 7A outlines the reaction conditions and probes used for labeling. Consistent with previous studies, it was essential to perform SPAAC on the azide present in the displayed protein of interest before conducting CuAAC to ensure that all free azides were consumed prior to exogenous addition of an azide probe for CuAAC. Flow cytometry analysis was used to evaluate all SPAAC and CuAAC reactions performed on doubly substituted proteins displayed on the yeast surface.

**Figure 7.**
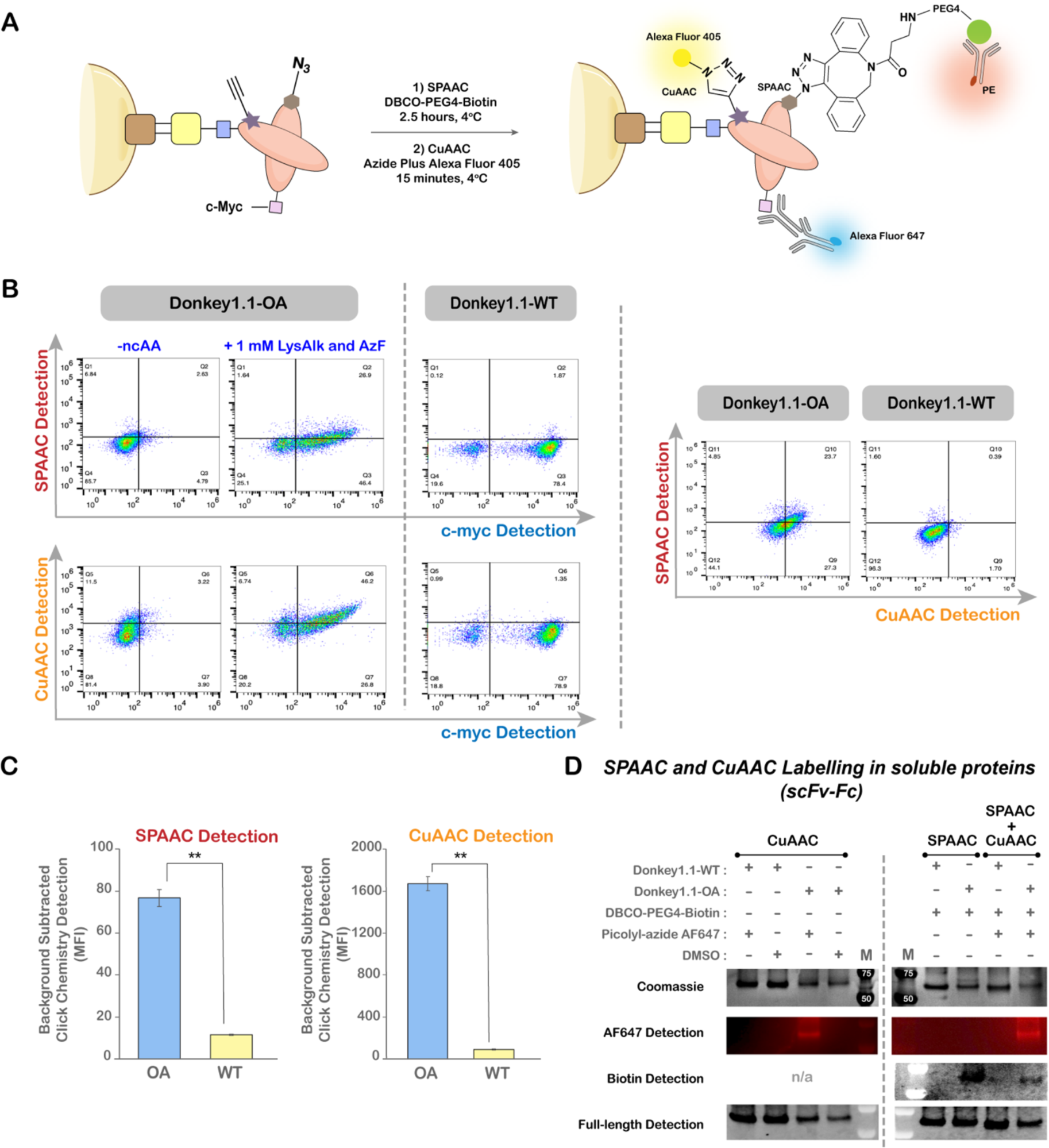
Site-specific double labeling of yeast-displayed and soluble proteins. A) Reaction and detection schemes for double labeling of yeast displayed, doubly substituted antibody constructs. TAG was employed to encode AzF (**4**; SPAAC reactions), while TGA was employed to encode LysAlk (**5**; CuAAC reactions). SPAAC products were detected using a PE-labeled anti-biotin antibody, CuAAC reaction products were detected with an Alexa Fluor 405-labeled azide probe, and c-myc (full length proteins) was detected using an anti-c-myc primary antibody followed by an Alexa Fluor 647-labeled secondary antibody. B) Flow cytometry dot plots of SPAAC and CuAAC reactions along with full-length (c-myc) display detection for Donkey1.1-OA under different induction conditions. Donkey1.1-WT (no ncAA substitutions) served as a negative control. C) Median fluorescence intensities of SPAAC and CuAAC reaction products in c-myc+ populations for Donkey1.1-OA and Donkey1.1-WT constructs. Error bars denote standard deviations obtained from three independent experiments. Unpaired t-tests were performed to evaluate statistical significance (**: p<0.005). D) Single and double labeling of soluble Donkey1.1-OA and Donkey1.1-WT scFv-Fcs. SPAAC was performed using a DBCO-PEG4-biotin probe and detected by western blot using Streptavidin 488. CuAAC was performed using Picolyl-azide Alexa Fluor 647 and detected by fluorescent gel imaging. Full-length protein expression was detected by western blot using goat anti-human IgG Fc Dy488. Coomassie staining was performed to verify sample purity. “M” indicates gel lane with molecular weight marker.

We first examined each reaction individually with both Donkey1.1-OA and Donkey1.1-AO constructs to identify efficient reaction conditions. Flow cytometry data from single-labeling studies with SPAAC and CuAAC reactions revealed double-positive populations for Donkey1.1-OA signifying that the full-length OA construct (c-myc+) had successfully undergone either SPAAC (biotin+) or CuAAC (Alexa Fluor 405+) reactions (Figure S18). On other hand, for the Donkey1.1-AO reporter protein, the flow cytometry data provided clear evidence for SPAAC, but not for CuAAC reactions, where high background fluorescence was observed. This made it difficult to delineate specific CuAAC labeling (Alexa Fluor 405+) of full-length proteins (c-myc+) from non-specific labeling of yeast cells (Figure S18). We attribute the lack of specific signal to low levels of LysAlk (**5**) at the H54 site, which is observed to be a less permissive site for TGA suppression. Thus, we proceeded to perform sequential double labeling with the Donkey1.1-OA construct as it unambiguously supported selective bioorthogonal reactions (SPAAC and CuAAC) (Figure 7A). Dot plots indicated elevated detection levels of both biotin (SPAAC) and Alexa Fluor 405 (CuAAC) probes in c-myc+ populations of Donkey1.1-OA samples. Such signals were undetected in the case of c-myc– cells of Donkey1.1-OA or in any cells of Donkey1.1-WT samples (Figure 7B, left panel). Additionally, dot plots of biotin detection versus Alexa Fluor 405 detection confirmed that both the SPAAC and CuAAC probes were present in the same population of Donkey1.1-OA cells (Figure 7B, right panels). Quantitative evaluation of the c-myc+ populations for the SPAAC (biotin) and CuAAC (Alexa Fluor 405) probes further substantiated statistically significant increases in the fluorescent signal of Donkey1.1-OA samples in comparison to Donkey1.1-WT samples (Figure 7C). All these data provide direct evidence of double-labeling of yeast-displayed proteins and confirm that both azide and alkyne functionalities encoded within the Donkey1.1-OA antibody construct can be selectively addressed using sequential SPAAC and CuAAC reactions.

To validate our observations of double labeling in yeast-display format, we used the soluble form of Donkey1.1-OA to conduct evaluations of SPAAC and CuAAC reactivities in solution via SDS-PAGE and Western blotting; donkey1.1-WT protein was utilized as a negative control. With individual SPAAC (probe: DBCO-PEG4-biotin) and CuAAC reactions (probe: picolyl azide Alexa Fluor 647) in solution, we detected protein labeling only with Donkey1.1-OA and not with the Donkey1.1-WT control (Figure 7D), revealing the presence of two noncanonical functionalities in Donkey1.1-OA and confirming the selectivity of individual SPAAC and CuAAC reactions in solution. Finally, we explored double labeling of Donkey1.1-OA in single pot reactions (Figure 7D). Clear signals for both probes were evident in the Donkey1.1-OA sample subjected to sequential bioorthogonal reactions. To achieve these signals, we treated samples with low concentrations of the SPAAC probe for extended reaction times to attempt to minimize interference of unreacted DBCO with the CuAAC probe (picolyl azide Alexa Fluor 647). Overall, for the first time, we have successfully demonstrated the feasibility of performing two distinct conjugations on yeast-prepared, doubly substituted proteins in both yeast display format and on secreted, purified proteins.

## Discussion

In this study, we have identified and systematically characterized OTSs, gene deletions, and stop codon contexts that support (and enhance) the incorporation of diverse ncAAs in response to opal (TGA) codons in yeast. Combining these systems with mutually orthogonal amber suppression machineries, we established dual ncAA incorporation in yeast for the first time with both intracellular and yeast-displayed proteins. The successful incorporation of two distinct ncAAs at pre-defined sites within a protein was further confirmed using ESI-MS and tandem mass spectrometry analysis. Furthermore, in both formats, we could exploit the encoded ncAAs for biorthogonal conjugation reactions to achieve double labeling, demonstrating the utility of our engineered system for presenting multiple, distinct functionalities within a single protein.

In our initial investigations of ncAA incorporation in response to TGA, we determined that variants from three distinct classes of OTSs in yeast – *Ec*TyrRS/tRNA^Tyr^, *Ec*LeuRS/tRNA^Leu^ and *Ma*PylRS/tRNA^Pyl^ – supported significant ncAA incorporation without any detectable readthrough from either orthogonal tRNACUAs or from endogenous aaRS-tRNA pairs. Such versatile decoding of TGA allowed us to identify two different OTS combinations that support dual ncAA incorporation in yeast: TyrRS/tRNACUA^Tyr^ + LeuRS/tRNAUCA^Leu^ and TyrRS/tRNACUA^Tyr^ + PylRS/tRNAUCA^Pyl^. Both combinations facilitated incorporation of two distinct ncAAs in proteins expressed intracellularly, and the TyrRS/tRNACUA^Tyr^ + LeuRS/tRNAUCA^Leu^ combination further facilitated dual ncAA incorporation in yeast display format. However, we did observe some evidence for aberrant production of full-length protein when the dual incorporation systems were induced in the presence of only a single ncAA (most notably with TyrRS/tRNACUA^Tyr^ + LeuRS/tRNAUCA^Leu^). This may be attributable to one or more of the following: 1) mischarging of tRNA by non-cognate aaRSs (for example, tRNACUA^Tyr^ by LeuRS); 2) recognition of a non-cognate amino acid substrate by an aaRS (for example, TyrRS recognizing AzMF, instead of its “cognate” AzF substrate); or 3) mischarging of a tRNA with a cAA (for example, tRNACUA^Tyr^ is charged with tryptophan rather than the ncAA of interest). The first possibility is a known phenomenon observed by Chatterjee and coworkers when using a combination of *E. coli* aaRSs in mammalian cells^17^. Mutating the tRNA anticodon sequence can result in the tRNA becoming a substrate of a non-cognate *E. coli* aaRS. In the future, such crosstalk could be mitigated by engineering or designing tRNAs to reduce such phenomena. The second possibility (noncognate ncAA charging) arises due to the polyspecific nature of many engineered aaRSs. This challenge could be avoided by utilizing aaRSs with non-overlapping sets of substrates. High-throughput engineering efforts from our lab and others have demonstrated the possibility of engineering highly specific aaRSs that discriminate against closely related ncAA substrates^13, 38, 78^. Lastly, the third possibility (canonical amino acid mischarging) is a known challenge associated with engineering aaRSs for genetic code expansion^38, 78–79^. While a portion of this challenge can be mitigated by using high ncAA concentrations during induction of protein synthesis, the discovery of additional high-performing aaRS variants that discriminate against the canonical amino acids will be important going forward. Dual ncAA incorporation in yeast will certainly benefit from ongoing efforts to expand the availability and performance of OTSs to access a broader palette of genetically encodable ncAAs with useful chemistries^10, 33, 34, 36–39^.

Identification of mutually orthogonal pairs of OTSs in yeast enabled us to prepare doubly substituted synthetic antibodies with unique reactive functionalities. Albeit with a decrease in affinity, these synthetic antibodies retained their binding function and facilitated efficient labelling with two bioorthogonal chemistries. Sequential SPAAC and CuAAC reactions led to doubly functionalized antibodies in both yeast display and soluble formats, which has not previously been achieved in yeast-based ncAA incorporation systems. Other promising biorthogonal conjugation chemistries for double labelling explorations on the yeast surface and in yeast-produced proteins include strain-promoted inverse electron-demand Diels-Alder cycloaddition (SPIEDAC) and chemoselective rapid azo-coupling reaction (CRACR)^11–13, 17, 80, 81^. Doubly labeled proteins on the yeast surface are expected to enable applications such as FRET to study protein conformation and dynamics, as evidenced by prior work in mammalian cells^16, 82–84^. Beyond bioconjugation strategies, proteins with dual ncAAs can support applications including intramolecular protein stapling, tethering strategies for small molecules or other drug modalities, and simultaneous presentation of two distinct chemistries such as crosslinking chemistries or posttranslational modifications^3, 12, 13, 18, 72, 85^. Thus, the emerging dual ncAA incorporation strategies in this work along with the expanding collection of OTSs supporting diverse ncAA incorporation in yeast will streamline the realization of these intricate protein manipulations in the near future.

Despite the successful implementation of dual ncAA incorporation in yeast, the systems described here exhibit relatively low efficiencies due to the lower readthrough efficiency of TGA compared to TAG by the *E. coli* OTSs. One likely explanation is that the anticodon change from CUA to UCA is not well-tolerated within the tRNAs, indicating that the tRNA identity elements play a role in determining TGA decoding efficiency^86^. Such inefficiencies from poor tRNA recognition can be alleviated either by enhancing the expression levels of these tRNAs to increase tRNA abundance, or by engineering suppressor tRNAs to improve tRNA recognition^35, 65, 87, 88^. In addition to tRNA properties, the nucleotide context surrounding opal codons also dictates ncAA incorporation efficiency. The effects of changing the context around an opal codon appear to be similar to the effects of changing the context around an amber codon. This is consistent with prior work from our own group and work from other groups investigating the role of nucleotide context in influencing ncAA incorporation efficiency (bacteria, mammalian cells and yeast)^56, 60–62, 67^. Besides engineering individual translation components, genetic and genomic engineering approaches offer a complementary set of tools for further enhancing dual ncAA incorporation efficiency in yeast. Single-gene deletion yeast strains *ppq1Δ* and *tpa1Δ* demonstrated significant improvements in ncAA incorporation in response to TGA across different OTSs, codon positions, and a range of ncAAs. The enhancements observed are consistent with the enhancements measured for ncAA incorporation in response to TAG in our previous work^56^. Our findings suggest that improving our fundamental understanding of eukaryotic protein biosynthesis may lead to further enhancements in genetic code manipulation in yeast (and other eukaryotes). Recently, our group reported a genome-wide screen to identify single-gene deletions that enhance ncAA incorporation efficiency in response to the amber (TAG) codon^66^. Out of the several dozen hits we identified, many of the genes had functions that are apparently unrelated to protein biosynthesis, indicating that there are numerous routes available for engineering enhanced dual ncAA incorporation systems. Finally, bold genome engineering strategies such as the construction of a synthetic yeast genome (Sc2.0 project), free up codons for the addition of ncAAs to the genetic code^26, 89, 90^. “Blank” codons in such synthetic genomes are expected to greatly enhance the efficiency of ncAA incorporation and should be compatible with the dual ncAA incorporation machinery described in this work.

## Conclusions

The yeast *S. cerevisiae* is one of the most important model organisms for elucidating fundamental biology and is a valuable chassis for protein engineering and synthetic biology. Empowering yeast with the ability to encode multiple ncAAs in individual proteins will lead to exciting biotechnological advances including new classes of protein-based therapeutics, biomaterials, artificial enzymes, and biosensors. More fundamentally, elucidating the processes that support and enhance dual ncAA incorporation systems is expected to lead to a deeper understanding of eukaryotic translation processes and the identification of unknown factors that can enhance genetic code expansion in yeast. In our current work, the tools and conditions identified are indispensable elements for realizing such applications and provide general insights that can be applied to the implementation of multiple ncAA incorporation systems in additional eukaryotic cells and organisms.

## Author Contributions

P.L. and J.A.V – Project conceptualization; P.L., M.S.M., B.R.L., R.A.S., and J.A.V. – experimental designs; P.L., M.S.M., B.R.L. – performed experiments and analyzed data; P.L. – wrote original manuscript draft; P.L., M.S.M., B.R.L., R.A.S., and J.A.V. – performed review and editing of the manuscript.

## Supporting Information

Experimental details, supplemental flow cytometry plots, fluorescence intensity graphs, RRE and MMF graphs, SDS-PAGE gel, MALDI mass spectrometry data, list of primers utilized for different plasmid construction, list of vector backbones for all the reporters and OTSs, list of yeast strains and plasmid combinations, DNA sequences of ‘Linker’ region present in dual fluorescent reporter construct and DNA sequence of scFv reporters with orthogonal codon incorporation sites.

## Conflicts of Interest

P.L. and J.A.V declare competing financial interests: A provisional patent application (Application No.-63/319,729) has been filed by Tufts University for the dual ncAA incorporation systems described in this work. T2RS5, a variant of *Ec*TyrRS, and LysAlkRS3, a variant of *Ec*LeuRS used in this study, are sequence variants disclosed in a patent application filed by Tufts University-U.S. Patent Application No. PCT/US22/29775.

## Acknowledgements

This research was supported by grants from the National Institute of General Medical Sciences of the National Institutes of Health (1R35GM133471 to J.A.V. and R01GM132422 to R.A.S.). The content of this work is solely the responsibility of the authors and does not necessarily represent the official views of the National Institutes of Health. The authors would also like to thank Heather Amoroso and Alla Leshinsky at the Koch Institute Biopolymers and Proteomics Core for their assistance with MALDI mass spectrometry data collection. Additionally, the authors would like to thank Rebecca Hershman for her valuable feedback on the manuscript.

## Notes

### Summary of Updates

We have added extensive mass spectrometry data to confirm the incorporation of two distinct ncAAs into the same biosynthesized protein in yeast. Statistical analysis of some experiments has been added. The manuscript has also been edited for clarity.

