## Supporting information for "Dual Noncanonical Amino Acid Incorporation Enabling Chemoselective Protein Modification at Two Distinct Sites in Yeast"

#### Contents:

1. Materials and Methods……S3
2. Figure S1……S14
3. Figure S2……S15
4. Figure S3……S16
5. Figure S4……S17
6. Figure S5……S18
7. Figure S6……S19
8. Figure S7……S20
9. Figure S8……S21
10. Figure S9……S22
11. Figure S10……S23
12. Figure S11……S24
13. Figure S12……S25
14. Figure S13……S26
15. Figure S14……S27
16. Figure S15……S29
17. Figure S16……S30
18. Figure S17……S31
19. Figure S18……S33
20. Table S1……S34
21. Table S2……S35
22. Table S3……S36
23. Table S4……S37
24. References……S38

### Materials and Methods

**Materials.**

Synthetic oligonucleotides were purchased from Genewiz. Restriction enzymes for cloning were purchased from New England Biolabs (NEB). All reagents used for PCR and Gibson Assembly were also purchased from NEB. Sanger DNA sequencing was carried out at Quintara Biosciences (Cambridge, MA). *E. coli* chemically competent cells were prepared using the Mix and Go! Transformation Kit (Catalog no. - T3001) from Zymo Research. Zymo Research Frozen-EZ Yeast Transformation II kit (Catalog no. – T2001) was used to prepare chemically-competent yeast cells and perform plasmid transformations. Plasmid purification from *E. coli* cells was performed using Epoch Life Science GenCatch Plasmid DNA Mini-Prep Kits. Ampicillin and Kanamycin used for culturing *E.coli* cells transformed with antibiotic-resistant plasmids were purchased from VWR Life Sciences in salt form, sodium and sulphate forms, respectively. Penicillin-Streptomycin (100X) for yeast culture propagation was obtained from Corning Inc. Noncanonical amino acids (ncAAs) were purchased from the following companies: i) *O*-methyl-L-tyrosine (OmeY): Chem-Impex International; ii) $N^{\varepsilon}$-Boc-L-Lysine (BocK): Chem-Impex International; iii) *p*-propargyloxyl-L-phenylalanine (OPG): Iris Biotech; iv) *p*-azido-L-phenylalanine (AzF): Chem-Impex International; v) $N^{\varepsilon}$-propargyloxycarbonyl-L-lysine (LysAlk): AstaTech; vi) 4-azidomethyl-L-phenylalanine (AzMF): SynChem. Bovine-serum albumin (BSA) used for preparation of Phosphate-Buffered Saline with 0.1% BSA and for protein purification were purchased from Fisher Scientific and Proliant Biologicals, respectively. Primary and secondary antibodies used for flow cytometry labeling were chicken anti-cMyc (Gallus Immunotech), mouse anti-HA (BioLegend), goat anti-chicken Alexa Fluor 647 (Thermo Fisher Scientific), goat anti-mouse Alexa Fluor 488 (Thermo Fisher Scientific) and PE anti-biotin (eBioscience). Chrompure Donkey IgG whole molecule as a Biotin-SP conjugate (Jackson ImmunoResearch) was used for binding titrations. To perform strain-promoted azide-alkyne cycloaddition (SPAAC) and copper-catalyzed azide-alkyne cycloaddition (CuAAC) reactions, the following reagents were purchased from Click Chemistry Tools: Biotin-PEG4-Dibenzocyclooctyne (DBCO) (Catalog No.- A105), Biotin-Azide (Catalog No.- 1265), Azide Plus Alexa Fluor 405 (Catalog No.- 1474), Picolyl Azide Alexa Fluor 647 (Catalog No.- 1300), and tris-hydroxypropyltriazolylmethylamine (THPTA) (Catalog No.- 1010). The following additional reagents used for CuAAC were purchased from Sigma Aldrich: (+)-Sodium L-ascorbate, aminoguanidine hydrochloride, copper sulfate pentahydrate, Ethylenediaminetetraacetic acid (EDTA), and Dimethyl sulfoxide (DMSO). 15 mL and 0.5 mL 30 kDa Molecular Weight Cut-Off (MWCO) centrifugal filter units for buffer exchange were purchased from Millipore Sigma. Protein A resin used for scFv-Fc purification was purchased from GenScript. For SDS-PAGE and western blot analysis, 4–12% Bis-Tris mini gels, SimplyBlue Safestain and iBlot™ Transfer Stacks were procured from Thermo Fisher Scientific. The following secondary antibodies used for western blots were purchased from Thermo Fisher Scientific: Strepavidin Alexa Fluor 488, and goat anti-human IgG Fc DyLight 488. The mass spectrometry-grade product Trypsin Gold was purchased from Promega. C18 Zip-Tip columns for MS sample preparation were purchased from Millipore Sigma. Trypsin-digested protein samples were sent to the Koch Institute Biopolymers and

Proteomics Core for Matrix-assisted laser desorption ionization (MALDI) mass spectrometry analysis.

**Yeast strain construction and media preparation.**

The construction of *S. cerevisiae* strain RJY100 used in this study has been previously described in Van Deventer *et al*[1]. BY4741 was purchased from Horizon Discovery, while single-gene deletions of BY4741 (*ppq1Δ* and *tpa1Δ*) were gifts from the Fuchs Lab at Tufts University.

Solid and liquid media used in this study were LB, SD-CAA, SG-CAA, SD-SCAA, SG-SCAA and YPG. Preparations of all solid and liquid media have previously been described in detail[2,3]. Selective SD-SCAA and SG-SCAA were prepared with amino acids dropped out of the media as follows: *i)* without Tryptophan (TRP), Leucine (LEU), or Uracil (URA) for RJY100 strains transformed with plasmids derived from pCTCON2 and pRS315; *ii)* without LEU or URA for BY4741 strains transformed with plasmids derived from pRS416 and pRS315. SD-CAA and SG-CAA media were used for the RJY100 strain transformed only with plasmids derived from pCTCON2. YPG media was used for large-scale induction of protein secretion to produce soluble proteins using RJY100. Luria Bertani (LB) medium supplemented with appropriate antibiotics was used to culture *E. coli* DH5α.Z1 cells.

**Noncanonical amino acid stock preparation.**

All noncanonical amino acids (ncAAs) used in this study were purchased as L-isomers with working stocks prepared at 50 mM concentration[2]. Briefly, the requisite amount of ncAA was weighed out and dissolved at 90% of the final volume with deionized water. When needed, 1 N NaOH was added dropwise to dissolve the ncAA. Additional deionized water was added until the final volume was reached, followed by sterile filtration through a 0.2 μm filter. All filtered ncAA solutions were stored at 4 °C and used within 1 week of their preparation.

**Plasmid construction.**

The primers used for cloning all of the new plasmid constructs in this work are highlighted in Table S1. Different plasmids and their *E. coli* and yeast selection markers are outlined in Table S2.

Construction of pRS315-KanRmod-LeuOmeRS/tRNA$_{CUA}^{Leu}$, pRS315-KanRmod-T2RS5/tRNA$_{CUA}^{Tyr}$, pRS315-KanRmod-MaPylRS/tRNA$_{CUA}^{Pyl}$ and pRS315-KanRmod-LysAlkRS3/tRNA$_{CUA}^{Leu}$ have been described elsewhere[3–5]. These plasmids were used as templates for the construction of the following new plasmids (where the tRNA anticodon was changed from CUA to UCA): pRS315-KanRmod-LeuOmeRS/tRNA$_{UCA}^{Leu}$, pRS315-KanRmod-T2RS5/tRNA$_{UCA}^{Tyr}$, pRS315-KanRmod-MaPylRS/tRNA$_{UCA}^{Pyl}$ and pRS315-KanRmod-LysAlkRS3/tRNA$_{UCA}^{Leu}$, respectively. The template plasmids, except for LysAlkRS3/tRNA$_{CUA}^{Leu}$, were digested overnight with NdeI and XmaI at 37 °C. Similarly, pRS315-KanRmod-LysAlkRS3/tRNA$_{CUA}$ was digested with NdeI and HindIII. For changing the tRNA anticodon sequence from CUA to UCA, mutagenic primers were used to amplify the tRNA. Two forward and reverse primer sets were employed to amplify the tRNA region of LeuOmeRS/tRNA$_{CUA}^{Leu}$, T2RS5/tRNA$_{CUA}^{Tyr}$, MaPylRS/tRNA$_{CUA}^{Pyl}$ and LysAlkRS3/tRNA$_{CUA}^{Leu}$ via PCR and to include the desired mutation. The two amplified inserts exhibited 30-bp overlap with each other and with the pRS315-vector backbone. The amplified inserts were then gel-extracted, purified, and subjected to overlap-extension (OE) PCR to generate the desired full-length tRNA insert. Following gel-extraction and purification of both insert and digested vector, a two-fragment Gibson Assembly was performed as well as a negative control in which the insert was omitted from the assembly. Gibson assembly reactions were incubated for 1.5 hours at 50 °C and then transformed directly into chemically competent *E.coli* DH5α.Z1 cells. The transformed cells were revived for an hour in SOC media at 37 °C under shaking conditions and then plated onto LB agar plates supplemented with 50 μg/mL kanamycin, followed by incubation at 37 °C for 16 hours. Single colonies were inoculated in 5 mL of LB broth supplemented with 50 μg/mL kanamycin and grown for 16 hours at 37 °C under shaking conditions. Plasmids were extracted from saturated cultures using an Epoch Plasmid DNA Mini-prep kit following the manufacturer's protocol. Plasmids were sent for Sanger Sequencing to verify the incorporation of the desired mutation.

Construction of pCTCON2-BYG (Addgene plasmid #158144), pCTCON2-BXG-TAG (Addgene plasmid # 158127), and pCTCON2-BXG-altTAG (Addgene plasmid #158132) have been reported previously[3,6]. Construction of pCTCON2-BXG-TGA and pCTCON2-BXG-altTGA was achieved by using pCTCON2-BXG-TAG and pCTCON2-BXG-altTAG, respectively, as templates. All of the cloning steps and conditions for these new constructs were exactly same as outlined for cloning tRNA$_{UCA}$s in the former paragraph with minor modifications. Briefly, the TAG to TGA mutation was inserted using mutagenic primers by PCR, following which, the amplified inserts were then gel-extracted, purified and spliced together using OE PCR. The templates were digested overnight with EcoRI and PstI HF. Following gel-extraction and purification of both spliced insert and digested vector, a two-fragment Gibson Assembly was performed and then transformed into competent *E. coli* cells, followed by plating on LB agar supplemented with 50 μg/mL ampicillin. The obtained clones were sequence verified.

Following a similar cloning strategy, pRS416-BXG-TGA and pRS416-BXG-Alt-TGA plasmids were generated using pRS416-BXG and pRS416-BXG-Alt-TAG as templates. The cloning of these templates, pRS416-BXG (Addgene plasmid # 158145) and pRS416-BXG-Alt-TAG (Addgene plasmid # 158146) has been reported previously[6]. The templates were digested sequentially with two restriction enzymes. First, they were digested with XmaI overnight and then subjected to a 65 °C incubation for 20 min to inactivate the enzyme. The salt concentration of the single-enzyme-digested mixture was then adjusted to 100 mM NaCl before incubation with the second restriction enzyme, BglII, overnight. The resulting digested vector was gel extracted and purified. Next, the spliced inserts containing the desired mutation were generated, first, by PCR amplification to obtain individual inserts and then joined using OE PCR. Employing the digested vector along with the purified spliced insert, a two-step Gibson Assembly reaction was set-up and then transformed into competent *E. coli* cells. The successfully transformed clones obtained on LB agar plates supplemented with ampicillin were sequence verified.

Construction of the single-plasmid systems (SPSs) pRS416-BYG-TyrOmeRS, pRS416-BXG-TyrOmeRS, and pRS416-BXG-Alt-TAG-TyrOmeRS has been previously described[7]. pRS416-BXG-TyrOmeRS and pRS416-BXG-Alt-TAG-TyrOmeRS were used as templates to introduce TGA into reporters to generate SPS reporter variants containing dual orthogonal codons, termed pRS416-BX$_2$G-OA-TyrOmeRS and pRS416-BX$_2$G-AO-TyrOmeRS, respectively. The cloning procedures used to add TGA codons into TAG-containing SPSs were identical to the procedures outlined for the construction of pRS416-BXG-TGA and pRS416-BXG-Alt-TGA.

Construction of the yeast-displayed, dual orthogonal codon SPSs, pCTCON2-Donkey1.1-AO-TyrAcFRS and pCTCON2-Donkey1.1-OA-TyrAcFRS was accomplished by first generating pCTCON2-Donkey1.1-AO and pCTCON2-Donkey1.1-OA constructs. To assemble these constructs, pCTCON2-Donkey1.1-L93TAG and pCTCON2-Donkey1.1-H54TAG were used as templates to allow for the insertion of TGA at amino acid position H54 or L93 of the synthetic antibody via PCR. This led to the generation of amplicons with two stop codons, Donkey1.1-AO and Donkey1.1-OA, respectively. pCTCON2-Donkey1.1-WT was used as a vector template for this cloning, which was linearized by digestion with PstI HF and XhoI to remove the encoded synthetic WT-antibody from the construct (the generation of pCTCON2-Donkey1.1-WT, pCTCON2-Donkey1.1-L93TAG and pCTCON2-Donkey1.1-H54TAG constructs have been reported elsewhere[8]). Following the cloning procedure of pCTCON2-BXG-TGA and pCTCON2-BXG-Alt-TGA, the two dual stop codon containing yeast-displayed constructs were generated and sequence verified. Second, pCTCON2-Donkey1.1-AO and pCTCON2-Donkey1.1-OA were used as templates for generating PCR amplicons for insertion into yeast-displayed SPSs. Specifically, pCTCON2-FABP2.3.6-TyrAcFRS was used as a backbone to insert the two new amplicons in place of the antibody fragment FABP2.3.6 (The construction of pCTCON2-FABP2.3.6-TyrAcFRS was previously described[7]). pCTCON2-FABP2.3.6-TyrAcFRS was digested in a two-step process, where it was first digested with XmaI overnight at 37 °C, followed by heat inactivation of XmaI at 65 °C for 20 min. Next, the plasmid was incubated overnight with EcoRI and NcoI at 37 °C. The resulting digested vector was gel extracted and purified. A cloning procedure identical to the procedure used for construction of pRS416-BX$_2$G-OA-TyrOmeRS (highlighted in the earlier paragraph) was also followed for generating the new yeast-displayed SPS clones, pCTCON2-Donkey1.1-AO-TyrAcFRS and pCTCON2-Donkey1.1-OA-TyrAcFRS. The successful clones were obtained by transforming into competent *E. coli* cells , followed by plating them on LB agar plates supplemented with ampicillin and lastly, sequence verifying using Sanger sequencing.

To make the secretion construct pCHA-Donkey1.1-OA-TyrAcFRS-TAA to enable preparation of a soluble, secreted protein with dual ncAA substitutions, we first constructed an SPS version of our previously reported secretion construct pCHA-Donkey1.1-H54TAG-TAA[8]. Towards this goal, pCTCON2-FABP2.3.6-TyrAcFRS was used as a template for amplifying inserts encoding the TyrAcFRS/$tRNA_{CUA}^{Tyr}$ OTS. The two generated inserts were spliced together by

OE PCR, followed by gel extraction and purification. The plasmid pCHA-Donkey1.1-H54TAG-TAA was digested downstream of the antibody encoding region using XhoI and SacI HF at 37 °C overnight. The remaining cloning strategy was identical to the strategy used for pRS416-BX$_2$G-OA-TyrOmeRS, resulting in generation of new construct pCHA-Donkey1.1-H54TAG-TyrAcFRS-TAA. Next, this construct was used to prepare two overlapping PCR amplicons introducing the TGA mutation at L93. In addition, pCHA-Donkey1.1-H54TAG-TyrAcFRS-TAA was digested overnight with EcoRI and XmaI at 37 °C to generate a linearized vector backbone. The remaining cloning steps as well as the transformation and sequencing procedures were performed identically to the procedures outlined for the construction of pCTCON2-BXG-TGA.

**Yeast transformation, propagation, and induction.**

Zymo-competent *S. cerevisiae* strains (RJY100, BY4741, *ppq1Δ*, and *tpa1Δ*) were transformed with different combinations of plasmids outlined in Table S3. Depending on the plasmid combinations and the type of yeast strain employed for transformation, cells were plated on solid SD-CAA media (for pCTCON2 or pRS416-backbone), SD-SCAA –Trp –Leu –Ura media (for pCTCON2- and pRS315- plasmid combinations) or SD-SCAA –Leu –Ura media (for pRS416- and pRS315- plasmid combinations) and incubated at 30 °C until colonies appeared (2-3 days).

For performing RRE and MMF experiments in biological triplicates, three independent transformants were separately inoculated into 5 mL of the appropriate liquid media (with amino acid dropouts dependent on the particular strain and combination of plasmids used) supplemented with 1X penicillin-streptomycin (100X stock) to prevent bacterial growth and grown to saturation at 30 °C over 2–3 days under shaking conditions (300 rpm) (Table S3). $OD_{600}$s of saturated cultures were measured and the desired volume corresponding to $OD_{600}$ = 1 was aliquoted into a culture tube. Aliquoted cultures were then pelleted and resuspended into 5 mL of appropriate liquid media. The diluted secondary cultures were incubated at 30 °C for 4–6 hours to allow cultures to grow to an $OD_{600}$ between 2 and 5. Once the cultures were within the desired range of $OD_{600}$ values, the desired volume corresponding to $OD_{600}$ = 1 was aliquoted, pelleted, and resuspended in 2 mL galactose-containing media, SG-CAA, SG-SCAA –Trp –Leu –Ura or SG-SCAA –Leu –Ura with penicillin-streptomycin. The 2 mL cultures were then induced with either one or two ncAAs at 1 mM final concentrations, depending on the experimental design, and incubated at 20 °C for 16–18 hours under shaking conditions (300 rpm). Cultures without any ncAAs were also induced and incubated alongside the above cultures to be used as controls.

In some cases, saturated cultures were stored at 4 °C for up to 6 weeks and then revived. To initiate revival, refrigerated cultures were resuspended, and then 500 µL of each culture was aliquoted and pelleted to remove old media. The pellet was then resuspended in 5 mL fresh media (with appropriate amino acid dropouts), supplemented with 1X penicillin-streptomycin, and grown to saturation for 16-18 hours at 30 °C with shaking. The resulting saturated cultures were then prepared for induction as described above.

**Flow cytometry sample preparation and data collection.**

Following induction, the $OD_{600}$s of the samples were measured and used to aliquot 2 x $10^6$ cells to wells in 96-well V-bottom plates (assuming 1 mL of culture at $OD_{600}$ = 1 contains $1*10^7$ cells). The volume in each well was adjusted to 200 μL using 1X PBSA and centrifuged for 2 min at 2400 x *g* to pellet the cells. The supernatant was discarded, and the cell pellets were resuspended in 200 μL of 1X PBSA and centrifuged again to wash the cells. This washing step was repeated for a total of 3 times. For evaluation of expression levels and stop codon readthrough events using intracellular dual-fluorescent reporters (BFP-GFP derivatives), the pelleted cells were resuspended in 200 μL of 1X PBSA and subjected to analytical flow cytometry on an Attune NxT Flow Cytometer (Life Technologies). For data collection on each sample, the total number of events to be collected was set to 10,000 with a flow rate of 25 μL/min. The VL1 channel (height) was used to monitor N-terminal reporter (BFP) levels, and the BL1 channel (height) was used to monitor C-terminal reporter (GFP) levels.

For evaluating expression levels and stop codon readthrough events using yeast-displayed reporters, the washed samples were first labeled with 50 μL PBSA/well containing primary antibodies, where both mouse anti-HA and chicken anti-c-Myc antibodies were added at 1:500 dilutions. 96-well plates containing samples were incubated at room temperature for 30 min with shaking (150 rpm). After incubation, cells were washed thrice with ice-cold 1X PBSA and resuspended and subjected to labeling with 50 μL PBSA/well containing secondary antibodies, where both goat anti-mouse Alexa Fluor 488 and goat anti-chicken Alexa Fluor 647 were added at 1:500 dilution. The plate was incubated on ice for 15 min, washed twice with ice-cold, 1X PBSA, and resuspended in 200 μL PBSA for flow cytometry analysis. For data collection on each sample, the total number of events to be collected was set to 10,000 with a flow rate of 25 μL/min. Additionally, BL1 channel (height) was selected for Alexa Fluor 488 detection and RL1 channel (height) for Alexa Fluor 647 detection. Single color controls treated with only Alexa Fluor 488 or Alexa Fluor 647 reagents and an unlabeled control were analyzed to determine gating. For preparation of single-color controls, either any ncAA-induced sample or WT sample was considered. These samples were processed separately in 1.7 mL microcentrifuge tubes, following similar labeling and washing steps. The only modification was that the PBSA washing step was performed in 1000 μL instead of 200 μL

**Flow cytometry data analysis.**

FlowJo and Microsoft Excel were used for analysis of flow cytometry experiments. Detailed protocols for Median Fluorescence Intensity (MFI), RRE and MMF determination from flow cytometry data using dual-fluorescent reporters and yeast-displayed reporters have been described previously[9]. Briefly, we used FlowJo to gate out single cells. A polygonal gate was drawn on unlabeled (for yeast-displayed proteins) or uninduced (intracellular dual-fluorescent reporter proteins) samples on a log-scale plot of side-scatter area versus forward scatter area. Subsequently, this population was analyzed on log-scale plot of forward scatter height versus forward scatter width to further distinguish the single cell population from doublets and larger cell aggregates by drawing a second gate. Isolated single cell populations were then analyzed on a dot plot using the fluorescent channels corresponding to N-terminal and C-terminal fluorescences (height). A quadrant was placed onto this plot such that the bottom left quadrant (Q4) contained at least 95% of the unlabeled or autofluorescent cell population. Next, a histogram plot of N-terminal fluorescence detection was prepared, where the population exhibiting fluorescence above the background levels was gated as the 'N-terminal+' population. This 'N-terminal+' population included cells expressing truncated proteins, full-length proteins, or a mixture of the two. These gates were applied to all samples to be analyzed for a given experiment. Lastly, a table was exported to Microsoft Excel containing the following information: MFI of N-terminus detection on the N-terminus+ population; MFI of C-terminus detection on the N-terminus+ population; MFI of N-terminus detection of Q4; and MFI of C-terminus detection of Q4.

MFI values of full-length protein detection levels were obtained by subtracting MFI values of Q4 population (C-terminus detection) from MFI values of N-terminus+ population (C-terminus detection). Error bars shown in plots of MFI data represent the standard deviation of three biological replicates. RRE and MMF values were calculated using the equations outlined in the main text, Figure 1B. Exact methodologies for calculations of RRE, MMF, and propagated error have been previously described[2,6,9]

**Binding titrations on yeast surface.**

The protocol for binding titrations on yeast surface has been previously described[8]. Briefly, $1.5*10^6$ induced RJY100 cells displaying scFvs - Donkey1.1-WT, Donkey1.1-AO or Donkey1.1-OA (both Donkey1.1-AO and Donkey1.1-OA were induced under conditions leading to AzF and LysAlk ncAA incorporation) were aliquoted into 1.7 mL microcentrifuge tubes and washed thrice with 1 mL 1X PBSA. The pelleted cells were resuspended in 1 mL of 1X PBSA, out of which 300 μL of resuspended cells was removed and mixed with 0.3 μL of chicken anti-cMyc by vortexing gently. Biotinylated Donkey-IgG in 1X PBSA was serially diluted on 96-well V-bottom plates, starting at 1 μM with seven subsequent 4-fold serial dilutions and a final PBSA blank (no IgG). To all these 9 wells containing antigen or PBSA (50 μL), 10 μL of induced cells mixed with chicken anti-cMyc were added and incubated overnight at 4 °C on an orbital shaker (150 rpm). The final number of cells per well was approximately 15,000. After incubation, the samples were washed three times with ice-cold PBSA and subjected to secondary labeling as described in 'Flow Cytometry Sample Preparation and Data Collection'. Goat anti-chicken Alexa Fluor 647 and Streptavidin Alexa Fluor 488 were used as secondary antibodies. After secondary labeling, the cells were washed and processed for flow cytometry. Flow cytometry data collection was performed as above with the exception that 3,000 events were collected per sample instead of 10,000 events. All titrations were performed in technical duplicates.

Analysis of the flow cytometry data for obtaining the binding affinities were executed according to the protocol outlined previously[8]. The cMyc-positive portions of single cell populations were gated on a histogram plot of C-terminus detection. MFI of biotinylated antigen detection on the cMyc+ population was subtracted from MFI of biotinylated antigen detection of Q4 population, to obtain background-corrected MFI values of biotinylated antigen detection. For each titration, these MFI values were normalized to the MFI value of the cells treated with the highest concentration of IgG, and the data was fitted to the 'One Site-- Specific binding' equation on Graphpad Prism Version 8.3.0, to obtain apparent $K_D$ values with 95% confidence intervals (Note: For cases, where apparent $K_D$ was greater than 2 μM, no absolute values were stated, rather >2 μM was used for description)

**Mass Spectrometry.**

Following tryptic digestion, peptide fragments were injected onto an AdvanceBio Peptide 2.7 μm column (2.1 x 150 mm, Agilent) and were eluted with a water:acetonitrile gradient mobile phase with 0.1% formic acid (0.400 mL/min; 95% - 5% over 19 min). An Agilent 6530 QTOF mass spectrometer was utilized in positive mode with a dual electrospray ionization (ESI) source. MS spectra were acquired using the following settings: ESI capillary voltage, 4000 V; fragmenter, 150 V; gas temperature, 325°C; gas rate, 12 L/min; nebulizer, 40 psig. Data was acquired at a rate = 5 spectra per second and scan range of 300 – 3000 m/z. MS/MS spectra were acquired using the following settings: ESI capillary voltage, 4000 V; fragmentor, 150 V; gas temperature, 325°C; gas rate 12 L/min; nebulizer, 40 psig. MS/MS was acquired at 2 spectra per second with a mass range of 300 – 3000 m/z, with stringency set to a medium isolation width, and charge of $z$=2–6. After identification, precursor ions were subjected to iterative rounds of collision induced dissociation in the collision chamber and subsequent mass identification. A ramped collision energy was used with a slope of 2.6 and offset of -4.8 as well as a slope of 3 and offset of 2.

Following acquisition, all MS/MS spectra were combined for display. Analysis was completed using multiple programs. Agilent MassHunter Bioconfirm (v. B.010.00) was used to identify and match tryptic fragments of the Donkey1.1-WT and Donkey1.1-OA proteins, and identify b and y ion fragmentation, as well as calculate the observed and predicted mass for each fragment. Full length precursor ion *m/z*'s were calculated using Bioconfirm; error tolerance was within 5 ppm. Mass error analysis was completed by matching observed b and y fragment ions with predicted fragmentations generated in Bioconfirm, and error is reported in ppm and Da. Qualitative Analysis (v. 10.00) was used to generate extracted ion chromatograms (EIC), extracted compound chromatograms (ECC), and total compound chromatograms (TCC). Figures created in Adobe Illustrator.

**Single and double labeling of yeast-displayed scFvs with SPAAC and CuAAC.**

$2*10^6$ induced RJY100 cells displaying either Donkey1.1-AO or Donkey1.1-OA (encoding AzF and LysAlk) were aliquoted into a 96-well V-bottom plate alongside uninduced RJY100 cells and induced RJY100 displaying Donkey 1.1-WT scFv as negative controls. Cells were washed three times with 1X PBSA and then subjected to Strain-promoted Azide-Alkyne Cycloaddition (SPAAC) reactions. SPAAC labeling was carried out based on a previous protocol[8]. Briefly, cells were first resuspended in 199 μL PBS, pH 7.4. To these resuspended cells, 1 μL of DBCO-PEG4-Biotin probe (20 mM stock concentration, 100 μM final concentration) was added, and then cells were incubated at 4 °C on an orbital shaker (60 rpm) for 2.5 hrs. Following the incubation, the reaction mixture was centrifuged for 10 min at 3214 x *g* at 4 °C and the supernatant was discarded. The pelleted cells were washed three times with ice-cold 1X PBSA and subjected to Copper-catalyzed Azide-Alkyne Cycloaddition (CuAAC) reactions. CuAAC labeling was carried out based on previously described protocols with some modifications[8,10].

In brief, the following components were mixed together and used to resuspend pelleted cells: *i)* 221 μL PBS, pH-7.4, *ii)* 0.25 μL of 20 mM Azide Plus Alexa Fluor 405 (final concentration- 20 μM), *iii)* 3.8 μL of CuSO4:THPTA (1:2 ratio of 20 mM CuSO4:50 mM THPTA; pre-mixed), *iv)* 12.5 μL of 100 mM amino guanidine and *v)* 12.5 μL of 100 mM sodium ascorbate. Cells were then incubated at 4 °C for 15 mins under mild shaking conditions (60 rpm). For single labeling of yeast-displayed scFvs, the pelleted cells were subjected to either SPAAC or CuAAC reaction conditions as outlined above. Following completion of the reactions, cells were pelleted by centrifuging for 10 min at 3214 x *g* at 4 °C. Pelleted cells were then washed three times with ice-cold 1X PBSA and subjected to primary and secondary antibody labeling, according to the protocol outlined in 'Flow cytometry sample preparation and data collection'. For this experiment, the primary antibody used was chicken anti-c-Myc (1:500 dilution), while the secondary antibodies used were goat anti-chicken Alexa Fluor 647 (1:500 dilution) and PE anti-biotin (1:500 dilution). For data collection, 10,000 events were collected with a flow rate of 25 μL/min. Three channels (VL1 for detection of Alexa Fluor 405, BL2 for detection of PE, and RL1 for detection of Alexa Fluor 647; all heights) were monitored to evaluate full-length display (cMyc expression) and attachment of probes.

For data analysis, the protocol outlined in 'Flow Cytometry Data Analysis' was followed with some minor modifications. The isolated single cell population were analyzed on a dot plot, where the x-axis corresponded to c-Myc detection and y-axis corresponding to either PE (SPAAC detection) or Alexa Fluor 405 (CuAAC detection). A quadrant was placed on the plot such that the bottom left quadrant (Q4) contained at least 95% of the unlabeled or autofluorescent cell population. Next, a histogram plot of c-Myc detection for the isolated single cell population was plotted and used to gate out the c-Myc+ population. From this population, MFIs corresponding to PE and Alexa Fluor 405 were extracted along with MFIs of double-negative populations (Q4). The MFIs of the double-negative populations were subtracted from the MFIs of the corresponding c-Myc+ population to obtain background-subtracted MFI values.

**Protein purification of Donkey1.1-OA and SDS-PAGE analysis.**

Transformation of pCHA-Fc-Sup-Donkey1.1-OA-TyrAcFRS along with pRS315-KanRmod-LysAlkRS3/$tRNA_{CUA}$ into competent RJY100 cells were performed according to protocols outlined in 'Yeast transformation, propagation and induction.' A single transformant was inoculated into 5 mL SD-SCAA –Trp –Leu –Ura media supplemented with 1X penicillin-streptomycin at 30 °C for 3 days with shaking (300 rpm). The saturated culture (with $OD_{600}$ ~

10) was diluted into 45 mL SD-SCAA –Trp –Leu –Ura media supplemented with penicillin-streptomycin and grown for 24 hours at 30 °C with shaking. The resulting 50 mL saturated culture was further diluted into 450 mL SD-SCAA –Trp –Leu –Ura media supplemented with penicillin-streptomycin and grown for an additional 24 hours at 30 °C with shaking. The following day, 500 mL saturated culture was pelleted at 3214 x *g* for 25–30 min and resuspended in 1 L YPG media supplemented with 0.1% BSA, 1X Penicillin-Streptomycin. To the resuspended culture, AzF and LysAlk were added to 1 mM final concentration, and then the culture was incubated at 20 °C with shaking for 4 days.

After induction, the culture was pelleted at 3214 x *g* for 25–30 min and the supernatant was collected. To the supernatant, 10X PBS, pH 7.4 was added to adjust the final concentration of the solution to 1X PBS; this supernatant was then sterile filtered using a 0.2 μm filter. To prepare for protein purification, Bio-Rad protein purification columns and reservoirs were washed with 1X PBS, and then 2 mL Protein A slurry was added to each column (1 mL resin volume). The resin was equilibrated with 1X PBS, followed by addition of the supernatant. The supernatant was passed twice through the resin, followed by three washes with 10 mL of 1X PBS each time. Finally, the protein was eluted using 7 mL of 100 mM glycine, pH 3.0, into a tube with 700 μL of 1 M Tris pH 8.5. The eluted fractions were stored at 4 °C overnight and buffer-exchanged either into water or 1X PBS using 15 mL centrifugal filter units with 30 kDa MWCO membranes.

Expression and purification of Donkey1.1-WT has been described previously and the same purified Donkey1.1-WT was employed in this study as well[5]. For SDS-PAGE analysis, 1-2 μg of protein was mixed with sterile water, 5 μL of 4X Bolt LDS sample, and 2 μL of 10× NuPAGE sample reducing agent and boiled at 100 °C for 5 min. The boiled samples were cooled at RT and the total sample volume (20 μL) was loaded onto 4–12% Bis-Tris mini gels along with Precision Plus Protein All Blue Prestained Protein Standard (Bio-Rad) and run for 22 min at 200 V. The gel was then washed three times in distilled water for 5 mins on an orbital shaker (60 rpm) and then stained for 1.5 hours in SimplyBlue SafeStain. Next, the gel was destained with distilled water overnight on an orbital shaker and imaged on an Azure c400 gel imager (Azure Biosystems). The remaining purified proteins were mixed with 100% glycerol to a final concentration of 50% v/v glycerol, flash frozen in liquid nitrogen and stored at –80 °C

**Tryptic digestions and sample preparation for MALDI-MS.**

Glycerol stocks of purified proteins were first buffer exchanged into sterile water using 0.5 mL centrifugal filter units with 30 kDa MWCO membranes. Next, 8 μg of each buffer-exchanged protein was boiled at 100 °C for 3 min and allowed to cool to RT before addition of 1 μg of mass-spectrometry grade trypsin. Mixtures were then incubated overnight at 37 °C and desalted using C18 Zip-Tip columns into 50% acetonitrile with 0.1% trifluoroacetic acid, following manufacturer's protocol. The resulting samples were then flash frozen and sent on dry ice to the Koch Institute Biopolymers and Proteomics Core for MALDI-MS.

**Single and double labeling of soluble proteins (scFv-Fcs).**

Prior to protein labeling, glycerol stocks of purified proteins were thawed and buffer exchanged into PBS using 0.5 mL centrifugation devices with 30 kDa MWCO membranes. Single labeling via either SPAAC or CuAAC with purified scFv-Fcs (Donkey1.1-OA and Donkey1.1-WT) was conducted in 50 μL reaction volumes. For SPAAC labeling, the reaction mixture was prepared in ice-cold 1X PBS, pH-7.4 by mixing 500 nM purified scFvs with 1 μM of DBCO-PEG4-Biotin. The reaction was allowed to proceed for 1.5 hours at 4 °C under mild mixing conditions (orbital rotor, 60 rpm), before quenching with *p*-azidophenylalanine (AzF) at a final concentration of 1 mM. For CuAAC labeling, the mixture was prepared in ice-cold 1X PBS, pH-7.4 by mixing together the following reaction components in the following order: i) 500 nM purified scFv; ii) 20 μM Picolyl azide dye (AF647 Picolyl Azide, Click Chemistry Tools); iii) 100 μM $CuSO_4$:200 μM THPTA (pre-mixed), iv) 5 mM Aminoguanidine and, v) 5 mM Sodium ascorbate. DMSO in place of the azide dye was used as a negative control. The reaction was allowed to proceed for an hour at 4 °C under mild mixing conditions (orbital shaker, 60 rpm). The reaction was then quenched with 2 μL of 100 mM EDTA.

For double labeling, the SPAAC and CuAAC conditions described above were employed, with SPAAC being executed first, followed by CuAAC. Importantly, no quenching step was performed in between the two reactions. However, the total reaction volume was maintained at 50 μL, with the amounts of SPAAC and CuAAC reagents adjusted accordingly. Following CuAAC, each of the individual quenching agents for SPAAC and CuAAC outlined above were added to the reactions.

Both single and double labeled protein samples were analyzed via SDS-PAGE and Western blotting. Triplicate sets of 4–12% Bis-Tris SDS-PAGE gels were run. One set of gels was imaged first on AF647 channel (Cy5) on an Azure Biosystems c400 imaging system for evaluating CuAAC reactions and then stained for the presence of proteins with Coomassie (SimplyBlue SafeStain). The other two gels were transferred to nitrocellulose membranes for Western blot, using an iBlot2 Dry Blotting System (Life Technologies). The nitrocellulose membranes were then blocked with 5% w/v BSA in Tris-Buffered Saline (TBS) + 0.1% v/v Tween20 (TBST) solution for an hour at room temperature under mild shaking conditions (orbital shaker, 60 rpm). The membranes were washed twice with 1X TBST for 10 min per wash at room temperature. One membrane was probed with Streptavidin Alexa Fluor 488 (1:1000 dilution in blocking buffer) to detect the presence of biotin (SPAAC reaction), and the other membrane was probed with Anti-Fc DyLight 488 (1:1000 dilution in blocking buffer) to detect the Fc portion of scFv-Fcs. The membranes were probed for an hour at 4 °C under mild shaking conditions (orbital shaker, 60 rpm) and then, washed thrice with 1X TBST (10 minutes per wash). Finally, the membranes were imaged using the AF488 channel (Cy2) on the c400 imaging system.

#### Supplementary Figures

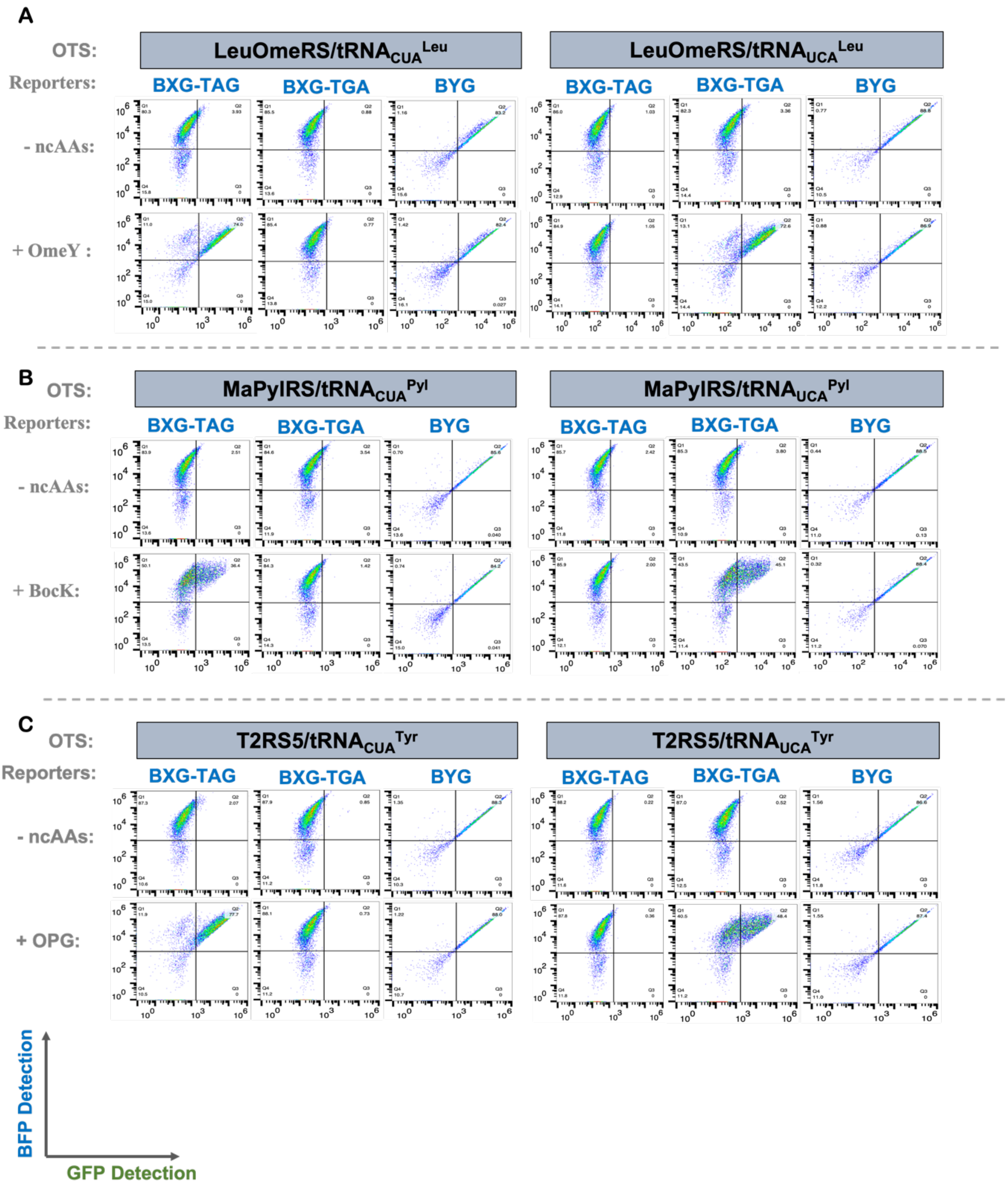


**Figure S1.** Flow cytometric plots depicting results of orthogonal codon readthrough BXG-TAG, BXG-TGA, and BYG reporters in the RJY100 yeast strain with A) LeuOmeRS/tRNA$^{Leu}$; B) MaPylRS/tRNA$^{Pyl}$; and C) T2RS5/tRNA$^{Tyr}$. Y-axis denotes N-terminal (i.e. BFP) detection and X-axis denotes C-terminal (i.e. GFP) detection of dual fluorescent reporters. Anticodon identity of tRNAs used in OTSs is noted in the figure. These plots are representative of biological triplicates performed for each sample and induction condition shown here.

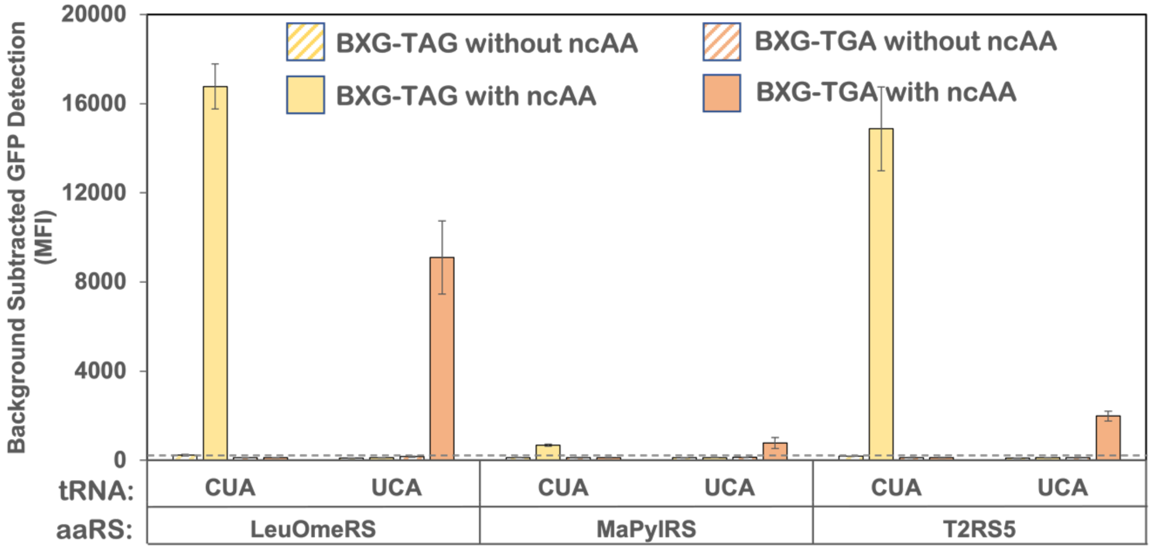


**Figure S2.** Background-subtracted Median Fluorescence Intensity (MFI) plots evaluating the expression of C-terminal reporter for BXG-TAG and BXG-TGA reporter constructs in the RJY100 yeast strain. Readthrough was evaluated with OTSs LeuOmeRS/tRNA$^{Leu}$, MaPylRS/tRNA$^{Pyl}$ and T2RS5/tRNA$^{Tyr}$ (anticodon identity noted in the figure). Cognate and non-cognate combinations of tRNA anticodon and reporter stop codon were evaluated for readthrough following induction in the presence or absence of an ncAA (*O*-methyl-L-tyrosine (**1**; OmeY) was used as substrate for LeuOmeRS/tRNA$^{Leu}$, $N^{\varepsilon}$-Boc-L-Lysine (**2**; BocK) was used as a substrate for MaPylRS/tRNA$^{Pyl}$ and lastly, *p*-propargyloxyl-L-phenylalanine (**3**; OPG) was used as a substrate for T2RS5/tRNA$^{Tyr}$). MFI data was obtained from the same flow cytometry experiments as depicted in Figure S1. The bars denote the average values of biological triplicates and error bars denote the standard deviation.

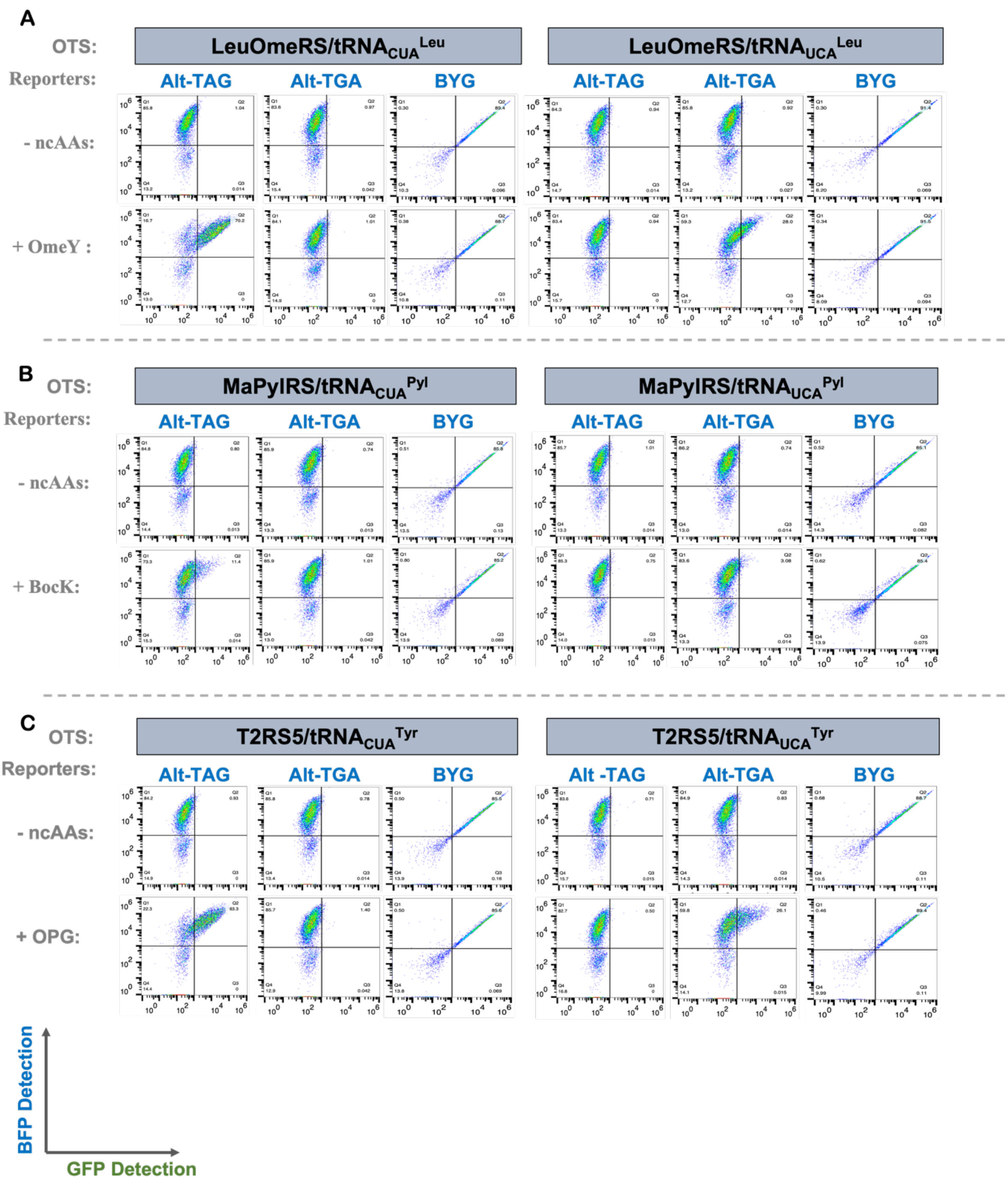


**Figure S3.** Flow cytometric plots depicting results of orthogonal codon readthrough of Alt-TAG, Alt-TGA, and BYG reporters in the RJY100 yeast strain with A) LeuOmeRS/tRNA$^{Leu}$; B) MaPylRS/tRNA$^{Pyl}$; and C) T2RS5/tRNA$^{Tyr}$ (anticodon identities shown in the figure). Y-axis denotes N-terminal (i.e. BFP) detection and X-axis denotes C-terminal (i.e. GFP) detection of dual fluorescent reporters. These plots are representative of biological triplicates performed for each sample and condition shown here.

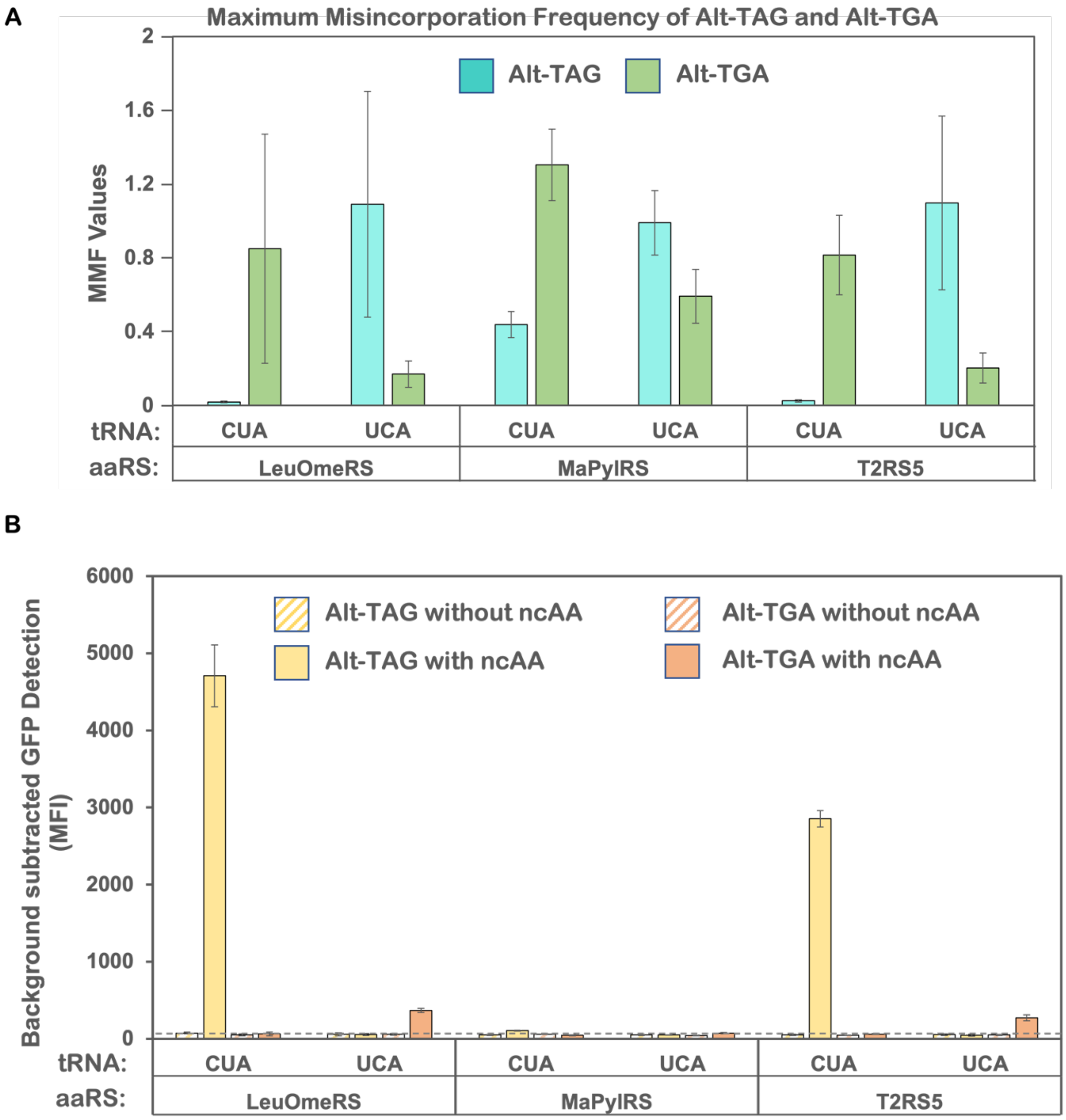


**Figure S4.** Quantitative evaluation of orthogonal codon readthrough with Alt-TAG and Alt-TGA reporter constructs in the RJY100 yeast strain. A) Maximum Misincorporation Frequency (MMF) of Alt-TAG and Alt-TGA reporters computed from RRE values with OTSs and tRNA anticodons as noted in the figure. The error bars are representative of three independent experiments, calculated from the standard deviation values and processed through error propagation equations. B) Background-subtracted Median Fluorescence Intensity (MFI) plots evaluating the expression of C-terminal fluorescent proteins for Alt-TAG and Alt-TGA reporter constructs. Readthrough was evaluated with OTSs LeuOmeRS/tRNA$^{Leu}$, MaPylRS/tRNA$^{Pyl}$, and T2RS5/tRNA$^{Tyr}$ (anticodon identities of tRNAs noted in the figure). MFI data was obtained from the same experiments used to generate the dot plots shown in Figure S3. The bars denote the average values of biological triplicates and error bars denote the standard deviations.

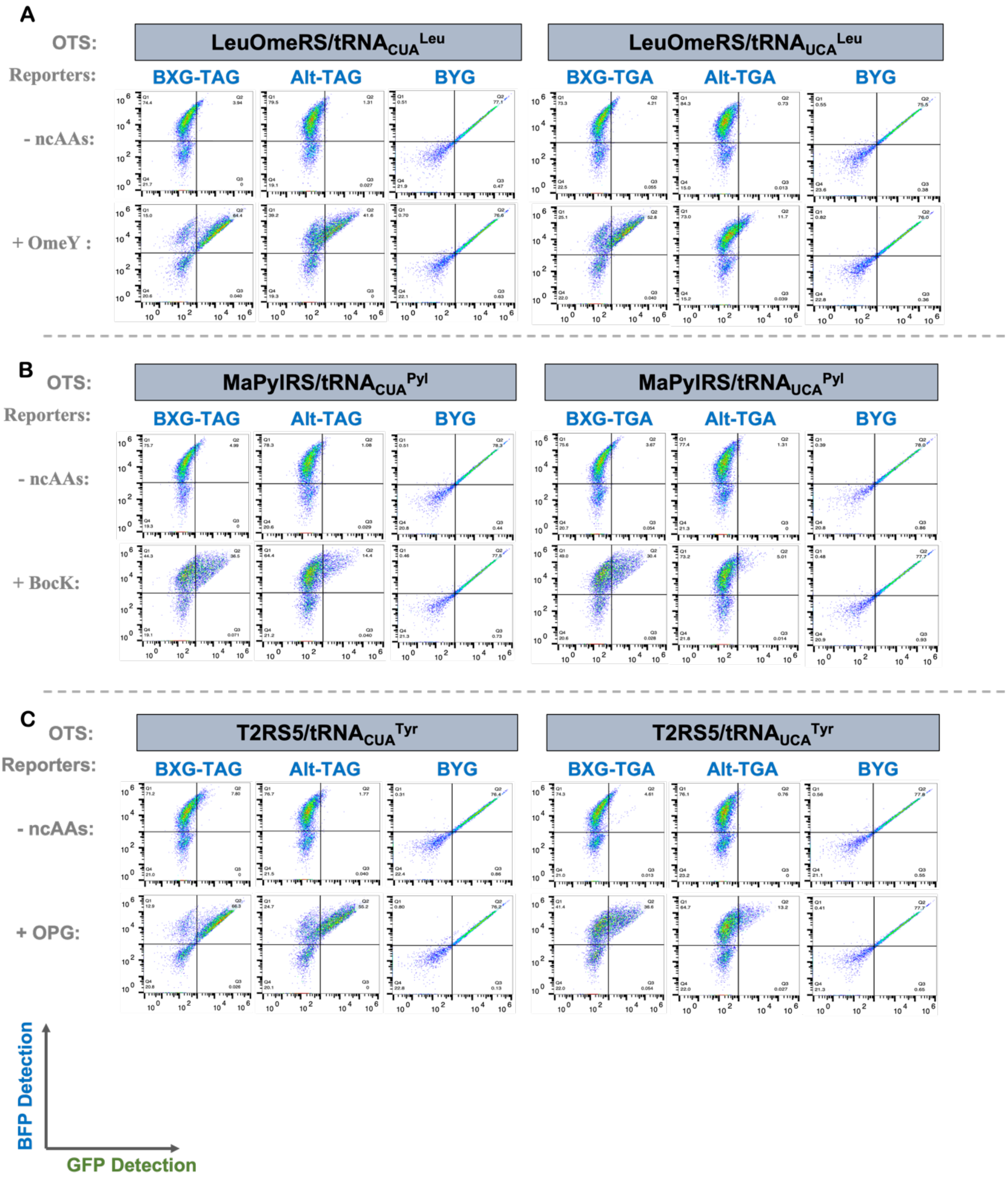


**Figure S5.** Flow cytometric plots depicting results of orthogonal codon readthrough of BXG-TAG, BXG-TGA, Alt-TAG, and Alt-TGA reporters with A) LeuOmeRS/tRNA$^{Leu}$; B) MaPylRS/tRNA$^{Pyl}$; and C) T2RS5/tRNA$^{Tyr}$ in the WT BY4741 yeast strain. The BYG reporter was employed as a positive control. Y-axis denotes N-terminal (i.e. BFP) detection and X-axis denotes C-terminal (i.e. GFP) detection of dual fluorescent reporters. These plots are representative of biological triplicates performed for each sample and condition shown here.

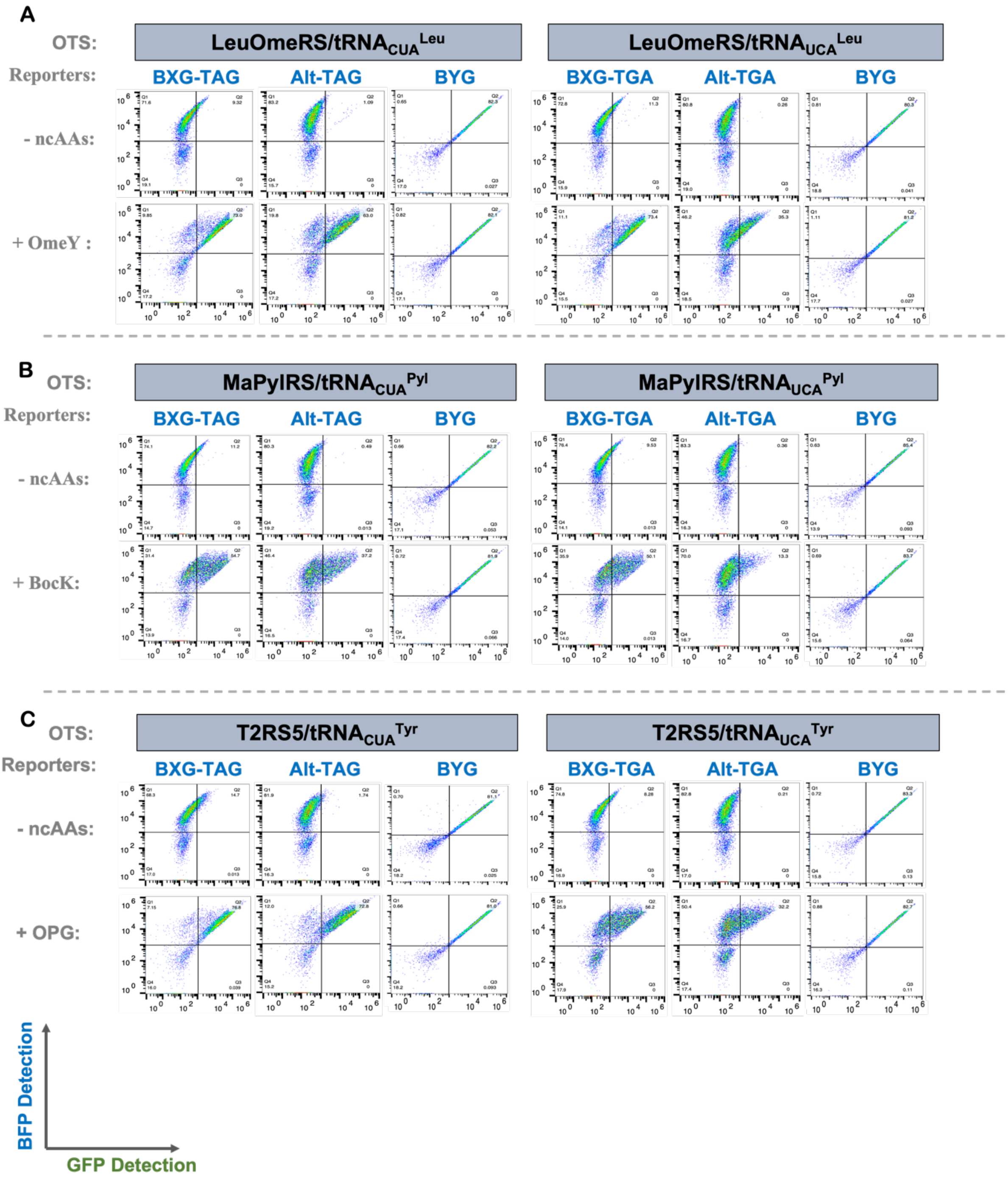


**Figure S6.** Flow cytometric plots depicting results of orthogonal codon readthrough of BXG-TAG, BXG-TGA, Alt-TAG, and Alt-TGA reporters with A) LeuOmeRS/$tRNA^{Leu}$; B) MaPylRS/$tRNA^{Pyl}$; and C) T2RS5/$tRNA^{Tyr}$ in the *ppq1Δ* yeast strain (anticodon identities of tRNAs noted in the figure). The BYG reporter was employed as a positive control. Y-axis denotes N-terminal (i.e. BFP) detection and X-axis denotes C-terminal (i.e. GFP) detection of dual fluorescent reporters. These plots are representative of biological triplicates performed for each sample and condition shown here.

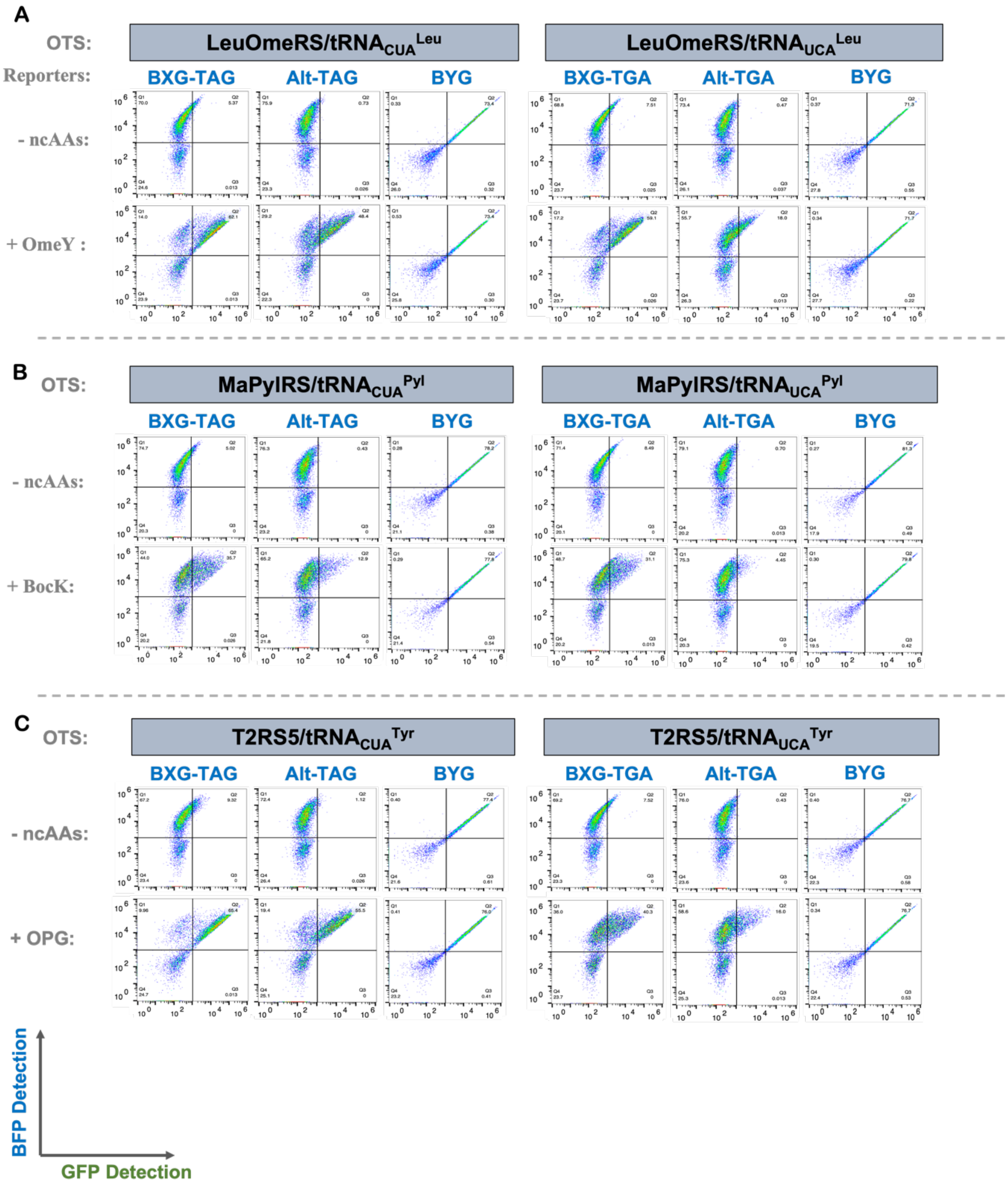


**Figure S7.** Flow cytometric plots depicting results of orthogonal codon readthrough of BXG-TAG, BXG-TGA, Alt-TAG, and Alt-TGA reporters with A) LeuOmeRS/$tRNA^{Leu}$; B) MaPylRS/$tRNA^{Pyl}$; and C) T2RS5/$tRNA^{Tyr}$ in the *tpa1Δ* yeast strain (anticodon identities of tRNAs noted in the figure). The BYG reporter was employed as a positive control. Y-axis denotes N-terminal (i.e. BFP) detection and X-axis denotes C-terminal (i.e. GFP) detection of dual fluorescent reporters. These plots are representative of biological triplicates performed for each sample and condition shown here.

A

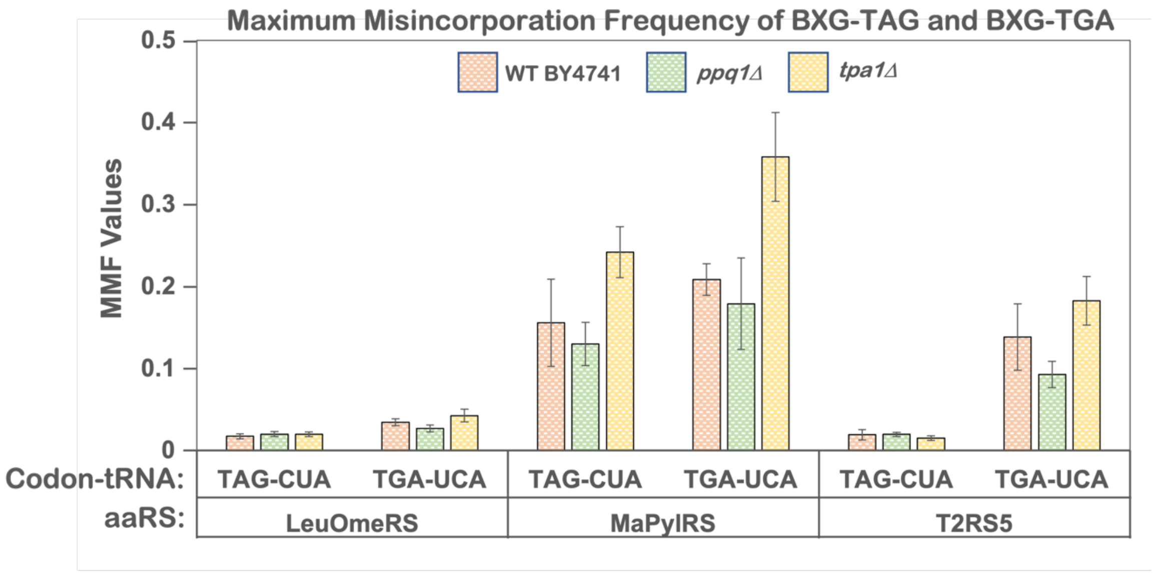


B

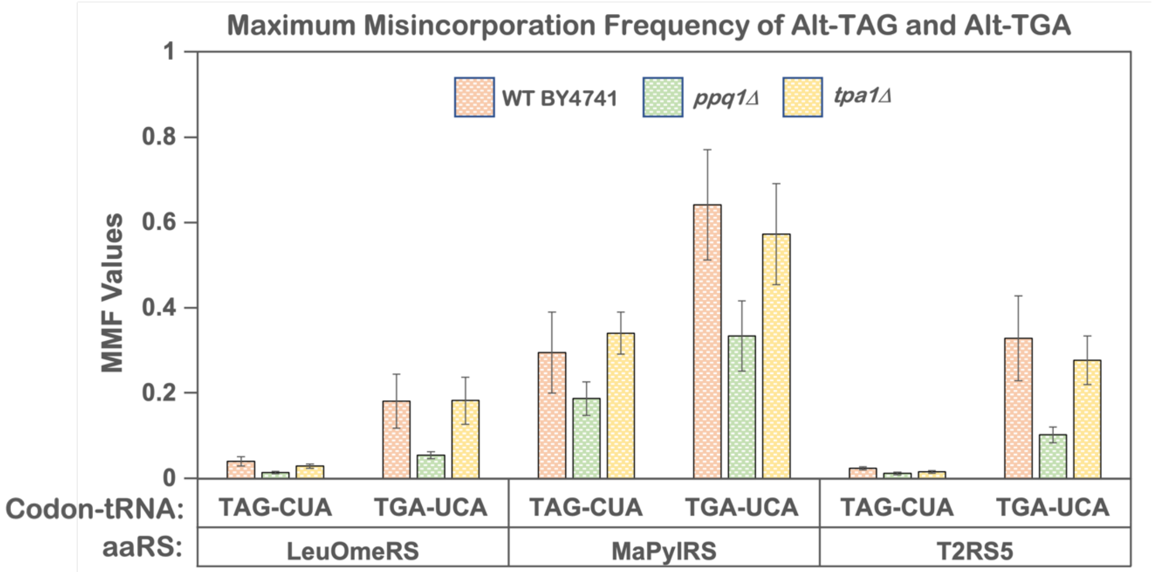


**Figure S8.** Maximum Misincorporation Frequency plots for the corresponding RRE data in main text Figure 3, for reporter constructs A) BXG-TAG and BXG-TGA; and B) Alt-TAG and Alt-TGA. Readthrough by each of the three OTSs encoding tRNAs with cognate anticodons in BY4741, *ppq1Δ*, and *tpa1Δ* strains. The error bars are representative of three independent experiments, calculated from the standard deviations and processed through error propagation equations.

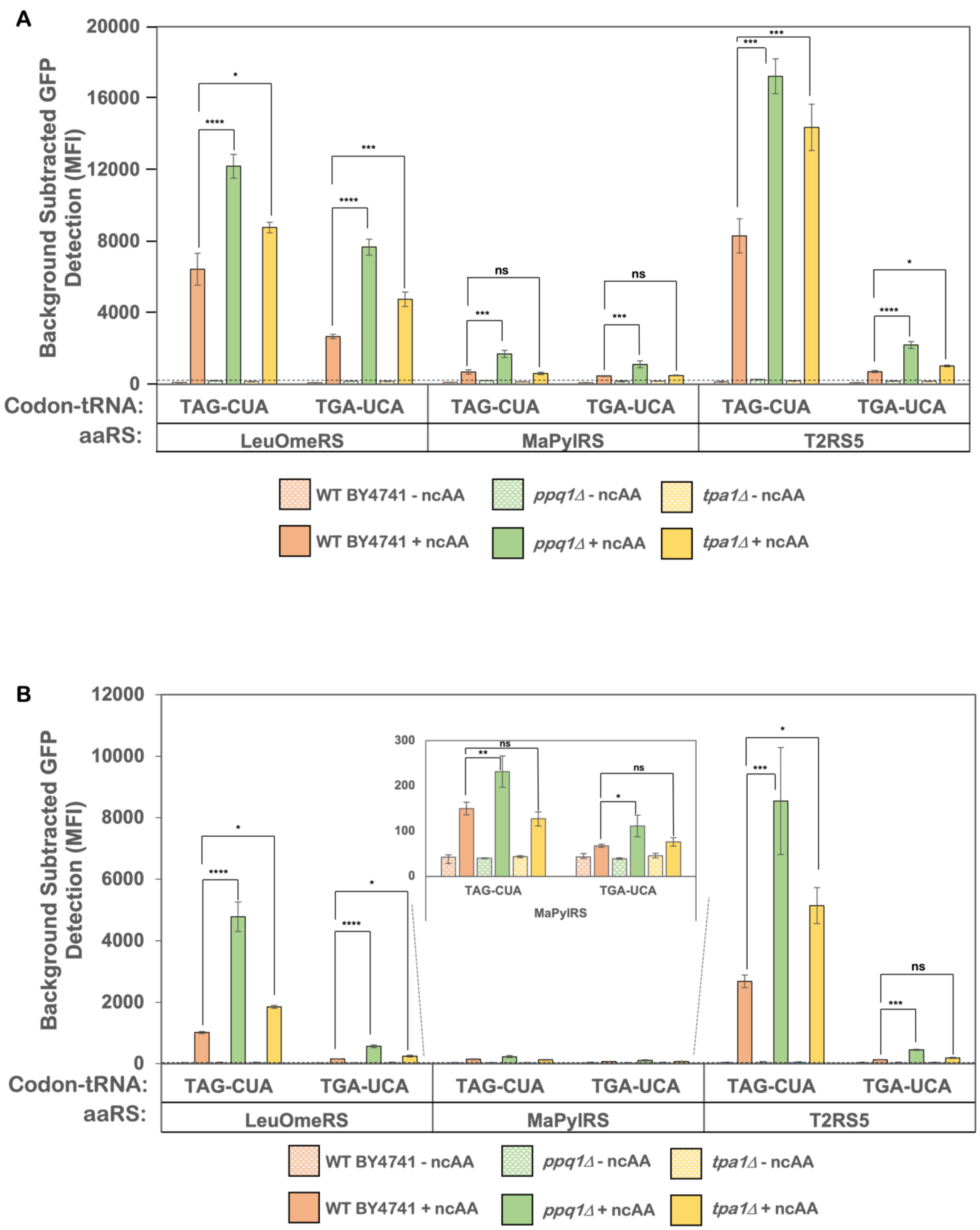


**Figure S9.** Background-subtracted Median Fluorescence Intensity (MFI) plots evaluating the expression of C-terminal fluorescent reporter protein for A) BXG-TAG and BXG-TGA; and B) Alt-TAG and Alt-TGA reporter constructs. Readthrough by each of the three OTSs encoding tRNAs with cognate anticodons in BY4741, *ppq1Δ*, and *tpa1Δ* strains. MFI data is obtained from the same experiments used to generate the dot plots shown in Figure S5-S7. The bars denote the average values of biological triplicates and error bars denote the standard deviations. One-way ANOVA was performed to determine the *p*-values between WT BY4741 + ncAA and knockout strains (*ppq1Δ*, and *tpa1Δ)* + ncAA samples. ns:non-significant; * - $p \leq 0.05$; ** - $p \leq 0.01$; *** - $p \leq 0.001$; **** - $p \leq 0.0001$.

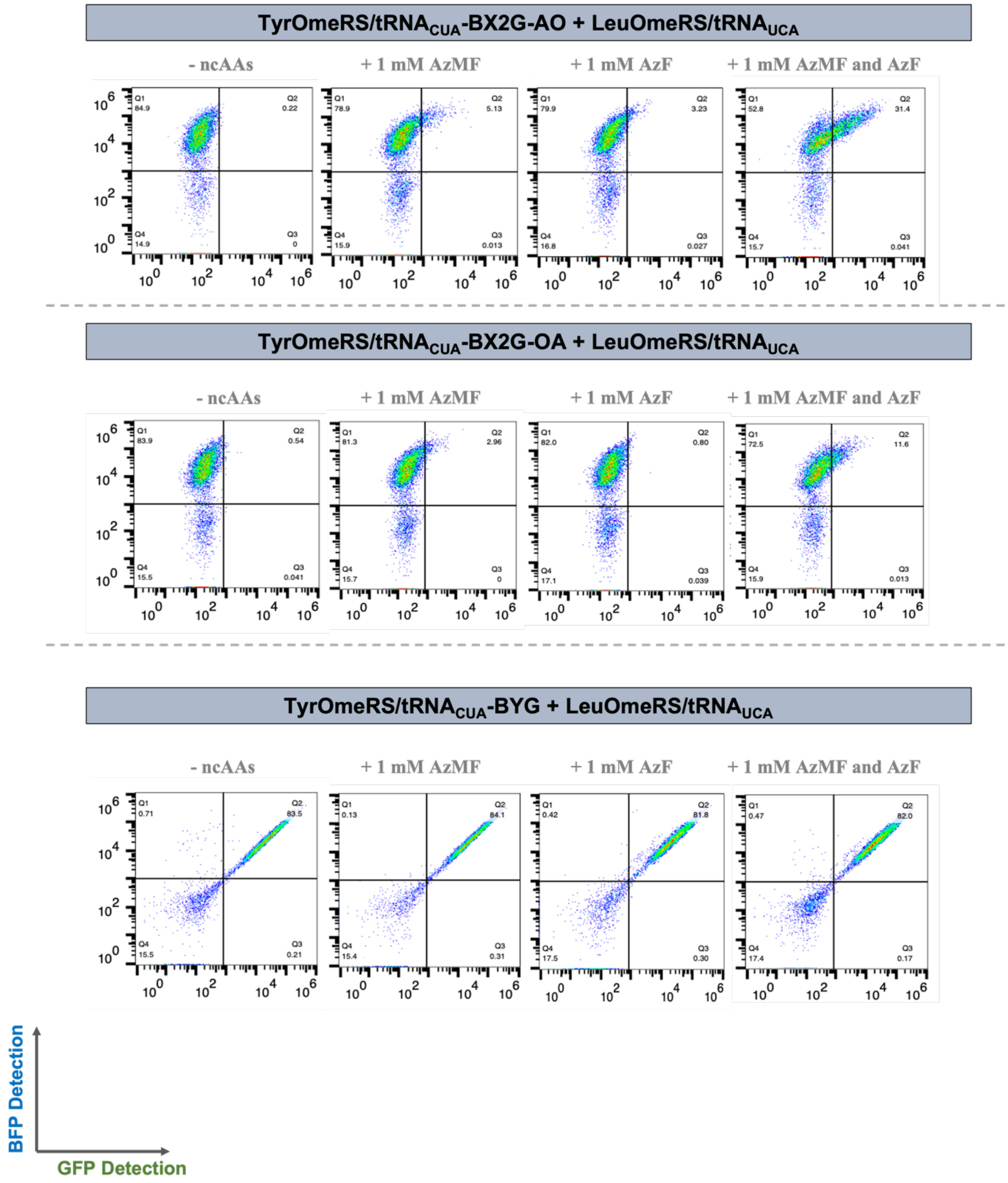


**Figure S10.** Flow cytometry dot plots of readthrough events of $BX_2G$-AO and $BX_2G$-OA in WT BY4741, employing the OTS combination *Ec*TyrRS/$tRNA_{CUA}^{Tyr}$ + *Ec*LeuRS/$tRNA_{UCA}^{Leu}$ and four different induction conditions in which 0, 1, or 2 ncAAs were added to the induction media. BYG was employed as a positive control. These dot plots are representative of experiments performed in biological triplicate for each sample and induction condition.

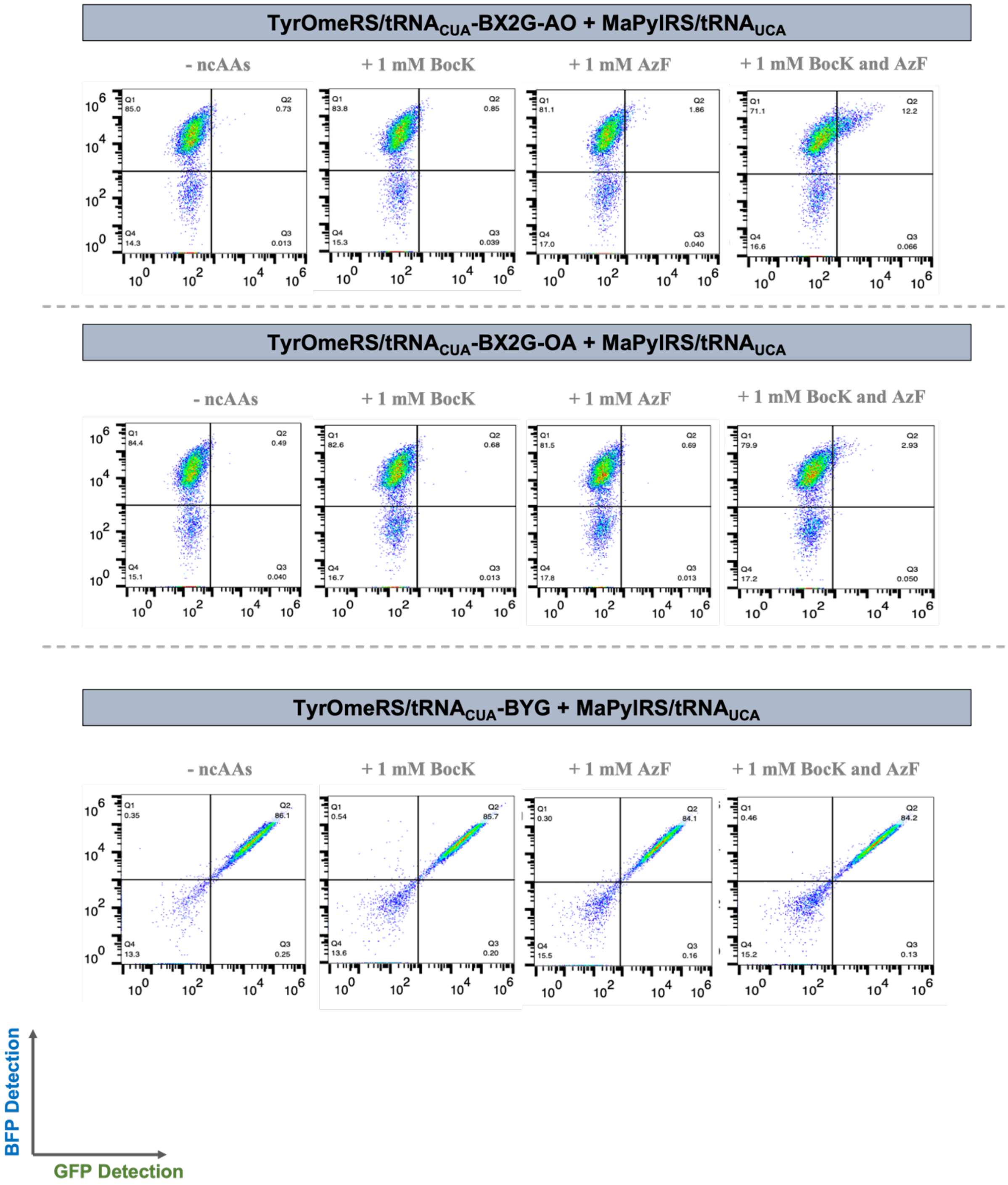


**Figure S11.** Flow cytometry dot plots of readthrough events of BX$_2$G-AO and BX$_2$G-OA in WT BY4741, employing the OTS combination *Ec*TyrRS/tRNA$_{CUA}^{Tyr}$ + *Ec*MaPylRS/tRNA$_{UCA}^{Pyl}$ and four different induction conditions in which 0, 1, or 2 ncAAs were added to the induction media. BYG was employed as a positive control. These dot plots are representative of experiments performed in biological triplicate for each sample and induction condition.

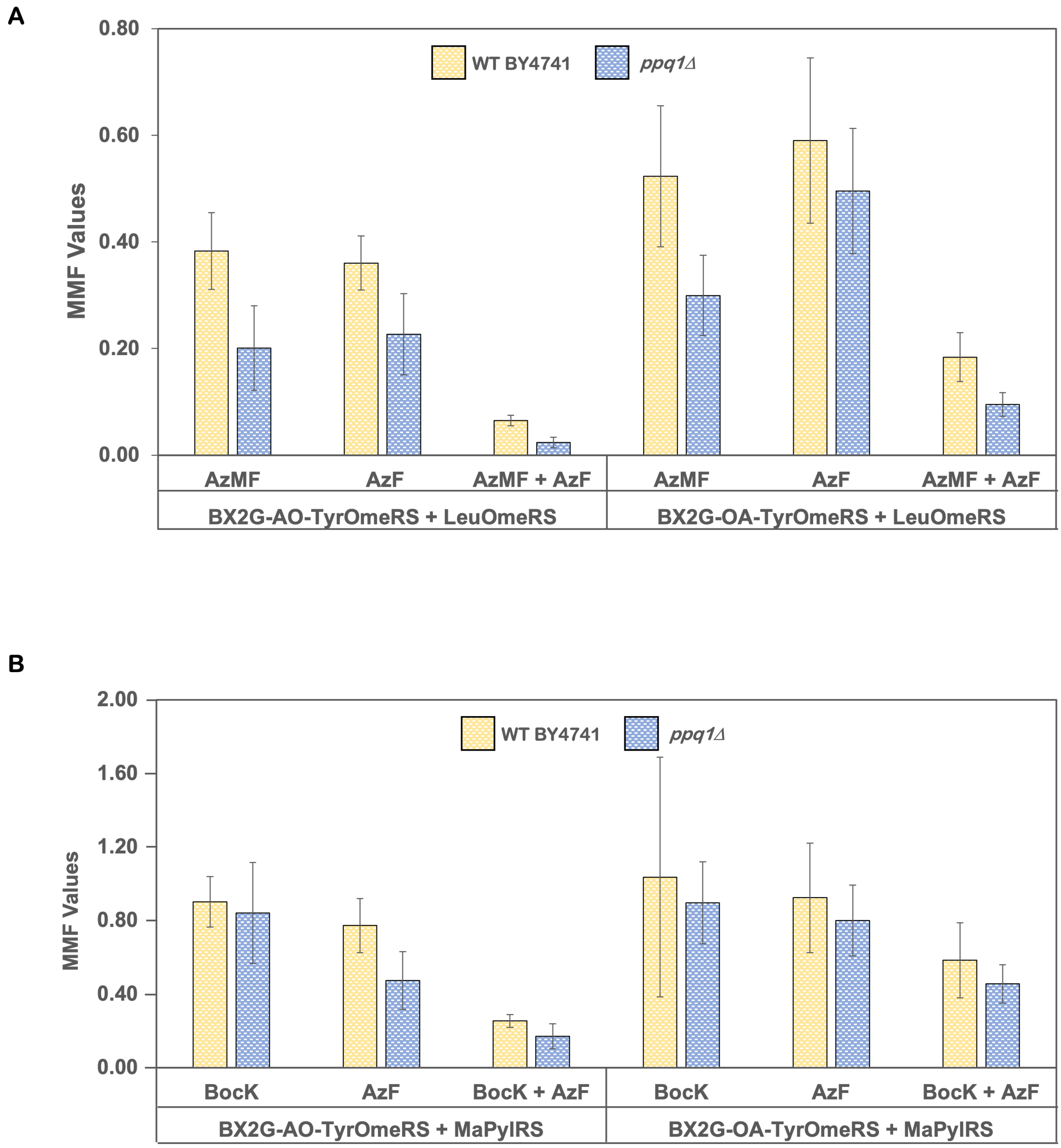


**Figure S12.** Maximum Misincorporation Frequency plots for the corresponding RRE data in main text Figure 4 using the reporter constructs BX2G-AO and BX2G-OA and OTS combinations of A) *Ec*TyrRS/tRNA$_{CUA}^{Tyr}$ + *Ec*LeuRS/tRNA$_{UCA}^{Leu}$; and B) *Ec*TyrRS/tRNA$_{CUA}^{Tyr}$ + *Ec*MaPylRS/tRNA$_{UCA}^{Pyl}$ in WT BY4741 and *ppq1Δ* yeast strains. The error bars are representative of three independent experiments, computed from the standard deviation values and processed through error propagation equations.

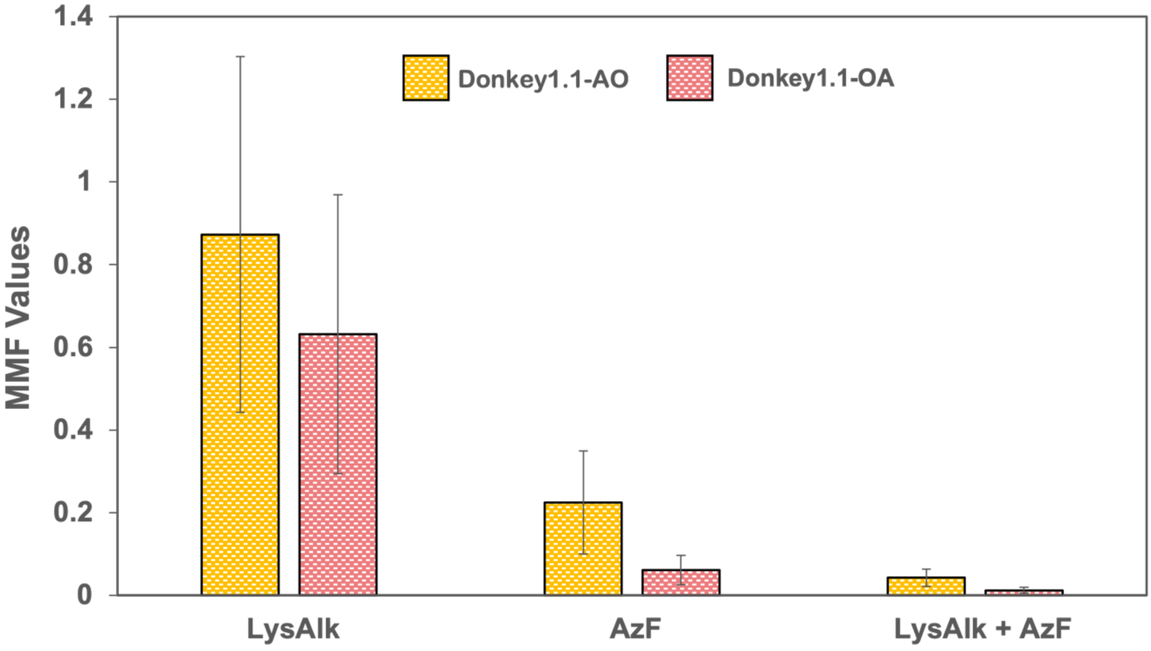


**Figure S13.** Maximum Misincorporation Frequency plots for the corresponding RRE data in main text Figure 5 using the reporter constructs Donkey1.1-AO and Donkey1.1-OA and the OTS combination of *Ec*TyrRS(TyrAcFRS)/$tRNA_{CUA}^{Tyr}$ + *Ec*LeuRS(LysAlkRS3)/$tRNA_{UCA}^{Leu}$ in the RJY100 strain. The error bars are representative of three independent experiments, calculated from the standard deviation values and processed through error propagation equations.

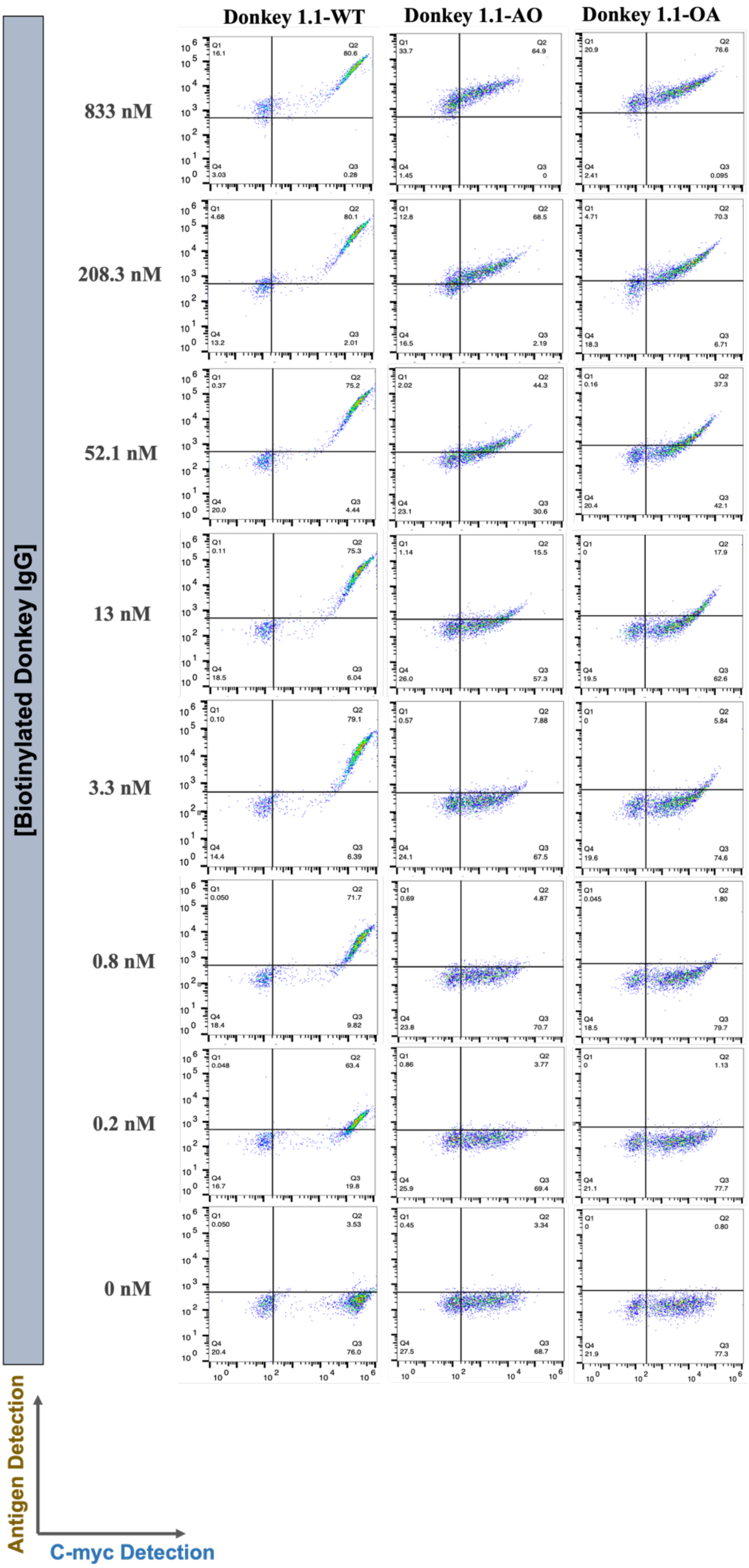


**Figure S14.** Flow cytometry dots plots of yeast display binding titrations of Donkey1.1-WT and its two dual ncAA-substituted variants, Donkey1.1-AO and Donkey1.1-OA, where the ncAAs are substituted at L93 and H54 positions of the scFv. Yeast-displayed clones were treated with varying concentrations of biotinylated Donkey IgG, starting from 1000 nM and titrating via 4-fold dilutions for a total of 7 concentrations. A negative control with no antigen was also included. Antigen binding was detected by labeling for biotin, while full-length expression of the reporter was detected by labeling for c-myc. The data obtained from these flow plots were analyzed to estimate binding affinities. For each clone, the experiment was performed in biological duplicates.

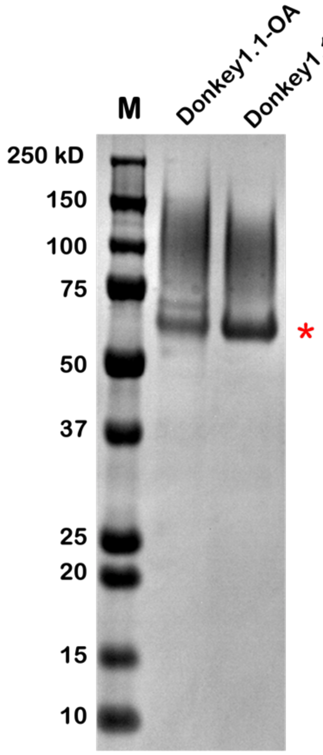


**Figure S15.** SDS-PAGE analysis of Protein A purified Donkey1.1-WT and Donkey 1.1-OA scFv-Fcs encoding both LysAlk and AzF. 4–12% Bis-Tricine gels were loaded under reducing conditions and stained with Coomassie SimplyBlue SafeStain.

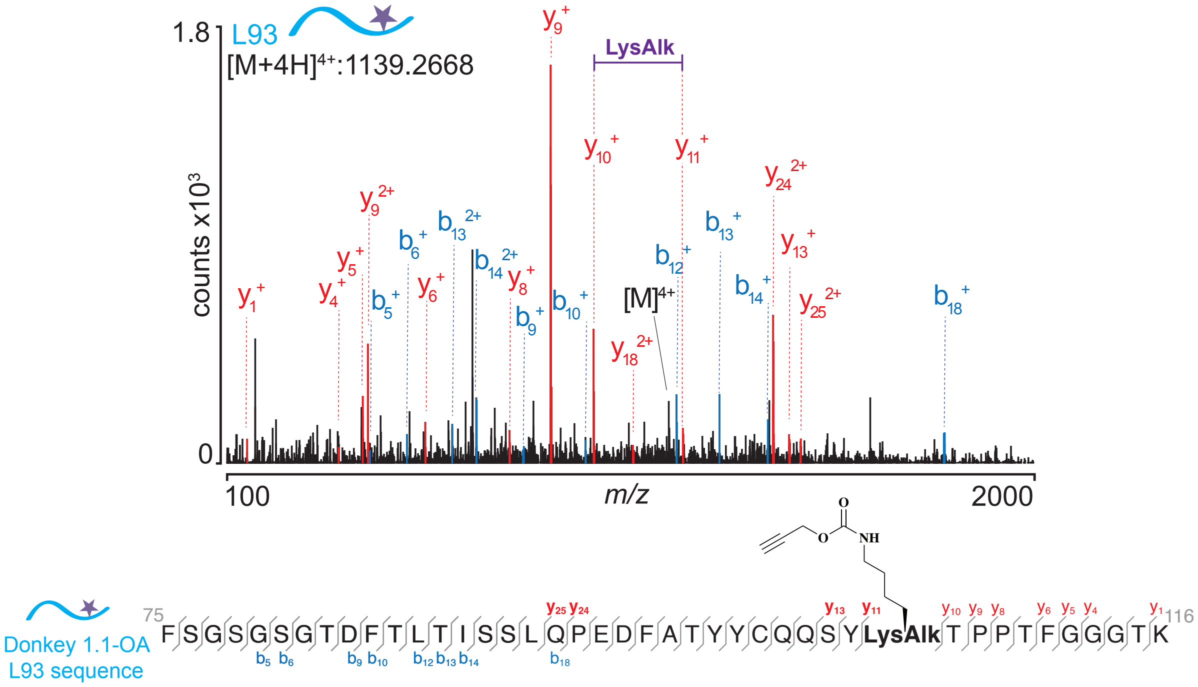


**L93 fragmentations**

| ion | z | $m/z_{expected}$ | $m/z_{observed}$ | error (Da) | error (ppm) |
|---|---|---|---|---|---|
| $b_5$ | 1 | 436.1779 | 436.1827 | -0.0048 | -10.9 |
| $b_6$ | 1 | 523.2084 | 523.2147 | -0.0063 | -12.1 |
| $b_9$ | 1 | 796.2982 | 796.3108 | -0.0126 | -15.9 |
| $b_{10}$ | 1 | 943.3833 | 943.3792 | 0.0041 | 4.4 |
| $b_{12}$ | 1 | 1157.4971 | 1157.5109 | -0.0138 | -12 |
| $b_{13}$ | 2 | 629.7906 | 629.783 | 0.0076 | 0.5 |
| $b_{14}$ | 2 | 686.3313 | 686.325 | 0.0063 | 17.4 |
| $b_{18}$ | 1 | 1786.824 | 1786.8494 | -0.0254 | -14.2 |

| ion | z | $m/z_{expected}$ | $m/z_{observed}$ | error (Da) | error (ppm) |
|---|---|---|---|---|---|
| $y_{25}$ | 2 | 1449.1199 | 1449.1438 | -0.0239 | -16.5 |
| $y_{24}$ | 2 | 1384.6096 | 1384.6129 | -0.0033 | -2.4 |
| $y_{18}$ | 2 | 1054.9610 | 1054.9768 | -0.0158 | -14.9 |
| $y_{13}$ | 1 | 1422.6878 | 1422.6009 | -0.0022 | -1.5 |
| $y_{11}$ | 1 | 1172.6082 | 1172.5946 | 0.0136 | 11.5 |
| $y_{10}$ | 1 | 962.4939 | 962.4942 | -0.0003 | -0.3 |
| $y_9$ | 2 | 861.4463 | 861.4465 | -0.0002 | -4.6 |
| $y_9$ | 1 | 431.2289 | 431.2269 | 0.0020 | 0.2 |
| $y_8$ | 1 | 764.3981 | 764.3937 | 0.0044 | 5.7 |
| $y_6$ | 1 | 566.2912 | 566.2933 | -0.0021 | -3.7 |
| $y_5$ | 1 | 419.2322 | 419.2249 | 0.0073 | 17.5 |
| $y_4$ | 1 | 362.1992 | 362.2034 | -0.0042 | -11.6 |
| $y_1$ | 1 | 147.1129 | 147.1128 | 0.0001 | 0.5 |

**Figure S16.** Complete mass list and error analysis for L93 peptide tandem mass spectrometry.

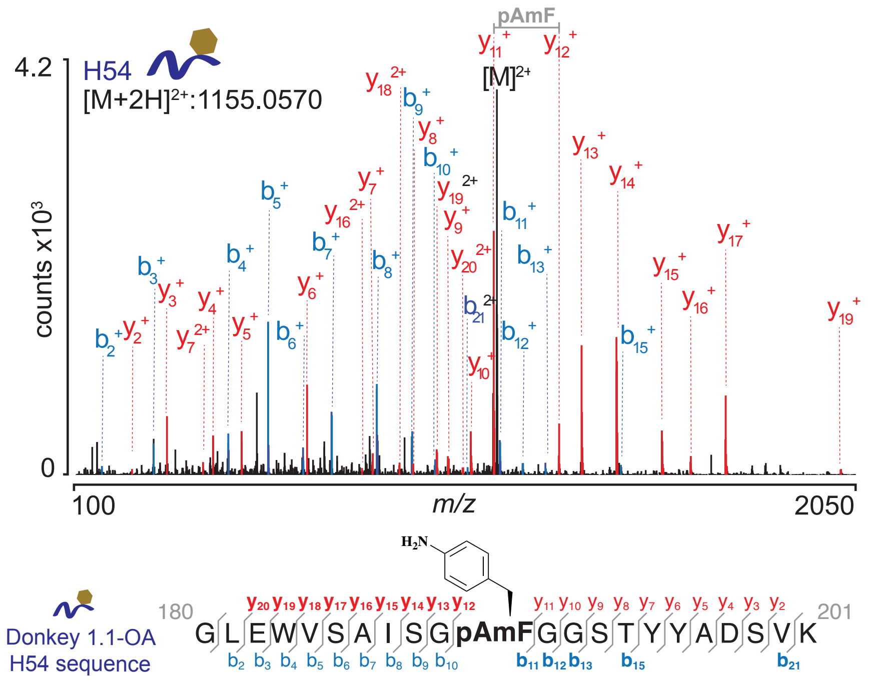


**H54 fragmentations**

| ion | z | $m/z_{expected}$ | $m/z_{observed}$ | error (Da) | error (ppm) |
|---|---|---|---|---|---|
| $b_2$ | 1 | 171.1161 | 171.1128 | 0.0033 | 19 |
| $b_3$ | 1 | 300.1563 | 300.1554 | 0.0009 | 2.9 |
| $b_4$ | 1 | 486.2342 | 486.2347 | -0.0005 | -1 |
| $b_5$ | 1 | 585.3019 | 585.3031 | -0.0012 | -2.1 |
| $b_6$ | 1 | 672.3331 | 672.3352 | -0.0021 | -3.1 |
| $b_7$ | 1 | 743.3627 | 743.3723 | -0.0096 | -12.9 |
| $b_8$ | 1 | 856.4551 | 856.4563 | -0.0012 | -1.4 |
| $b_9$ | 1 | 943.4861 | 943.4884 | -0.0274 | -2.4 |
| $b_{10}$ | 1 | 1000.504 | 1000.5098 | -0.0058 | -5.9 |
| $b_{11}$ | 1 | 1162.5974 | 1162.5891 | 0.0083 | 7.1 |
| $b_{12}$ | 1 | 1219.6084 | 1219.6106 | -0.0022 | -1.8 |
| $b_{13}$ | 1 | 1276.6085 | 1276.6321 | -0.0236 | -18.5 |
| $b_{15}$ | 1 | 1464.694 | 1464.7118 | -0.0178 | -12.1 |
| $b_{21}$ | 2 | 1082.4909 | 1082.5068 | -0.0159 | -14.7 |

| ion | z | $m/z_{expected}$ | $m/z_{observed}$ | error (Da) | error (ppm) |
|---|---|---|---|---|---|
| $y_{20}$ | 2 | 1069.9873 | 1070.0051 | -0.0178 | -16.6 |
| $y_{19}$ | 2 | 1005.4957 | 1005.5098 | -0.0141 | -11.8 |
| $y_{19}$ | 1 | 2010.9429 | 2010.9637 | -0.0208 | -10.4 |
| $y_{18}$ | 2 | 912.427 | 912.4442 | -0.0172 | -18.8 |
| $y_{17}$ | 1 | 1724.8146 | 1724.8126 | 0.0002 | 1.1 |
| $y_{16}$ | 1 | 1637.7607 | 1637.7806 | -0.0199 | -12.1 |
| $y_{16}$ | 2 | 819.3853 | 819.3939 | -0.0086 | -10.6 |
| $y_{15}$ | 1 | 1566.7318 | 1566.7435 | -0.0117 | -7.5 |
| $y_{14}$ | 1 | 1453.6585 | 1453.6594 | -0.0009 | -0.6 |
| $y_{13}$ | 1 | 1366.6263 | 1366.6274 | -0.0011 | -0.8 |
| $y_{12}$ | 1 | 1309.6059 | 1309.6059 | 0.0000 | 0 |
| $y_{11}$ | 1 | 1147.532 | 1147.5266 | 0.0054 | 4.7 |
| $y_{10}$ | 1 | 1090.4993 | 1090.5051 | -0.0004 | -5.4 |
| $y_9$ | 1 | 1033.476 | 1033.4837 | -0.0077 | -7.4 |
| $y_8$ | 1 | 946.4354 | 946.4516 | -0.0162 | -17.2 |
| $y_7$ | 2 | 423.2057 | 423.2056 | 0.0001 | 0.3 |
| $y_7$ | 1 | 845.4115 | 845.404 | 0.0075 | 9 |
| $y_6$ | 1 | 682.3429 | 682.3406 | 0.0023 | 3.3 |
| $y_5$ | 1 | 519.2778 | 519.2773 | 0.0005 | 0.9 |
| $y_4$ | 1 | 448.2394 | 448.2402 | -0.0008 | -1.8 |
| $y_3$ | 1 | 333.2134 | 333.2132 | 0.0002 | 0.4 |
| $y_2$ | 1 | 246.1771 | 246.1812 | -0.0041 | -16.6 |

**Figure S17.** Complete mass list and error analysis for H54 peptide tandem mass spectrometry.

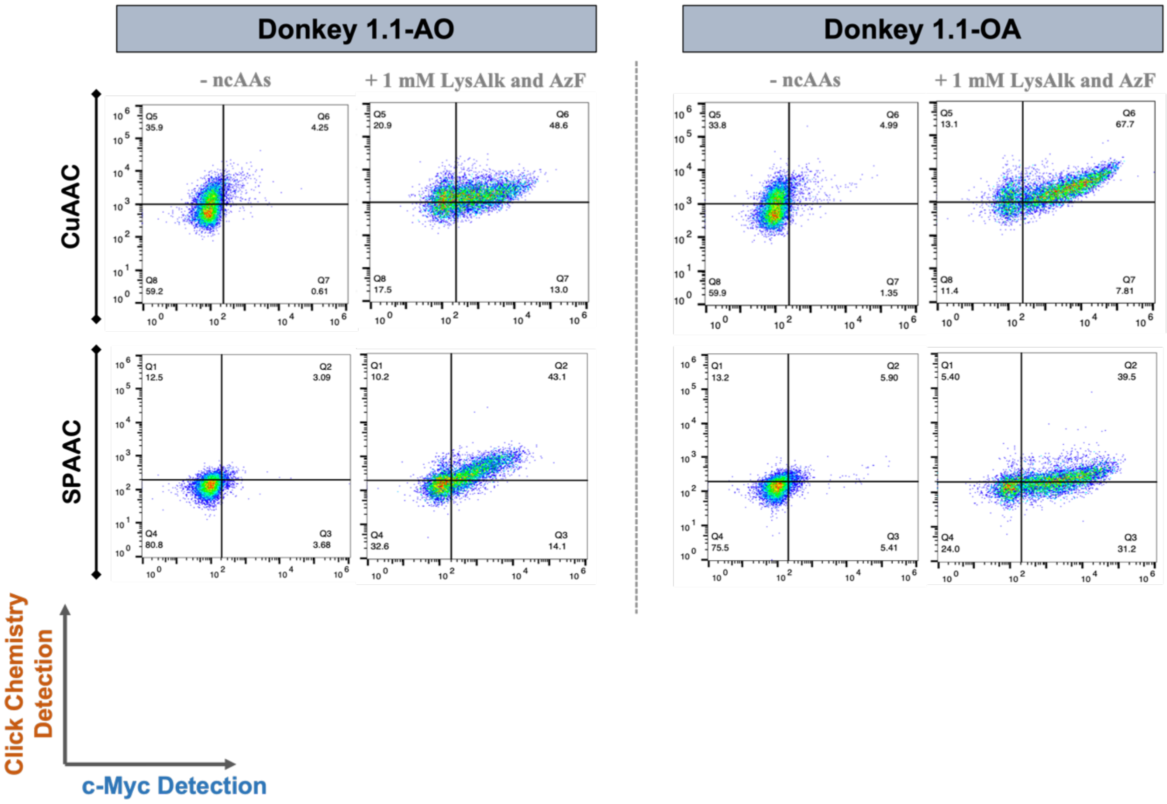


**Figure S18.** Flow cytometry evaluations of individual SPAAC and CuAAC labeling reactions with yeast displaying Donkey1.1-AO or Donkey1.1-OA after induction in the absence or presence of ncAAs. Y-axis denotes click chemistry probe detection and x-axis denotes full-length display detection (c-Myc). These plots are representative of biological triplicates performed for each sample.

#### Supplementary Tables

**Table S1.** Sequences of primers used for plasmid construction.

| No. | Primer Name | Primer Sequence (5'→3')* |
|---|---|---|
| #1 | pCTCON2-BFP-Fwd | TTAACGTCAAGGAGAAAAAACCCCGGATCG |
| #2 | pCTCON2_Rev_TGA_BFP_Linker | GCCCTTGGATGC**TCA**TTCGCCAGAACCAGCAGCGGAGCCAGCGGATCCCATATTAAGCTT |
| #3 | pCTCON2_Fwd_TGA_BFP_Linker | AAGCTTAATATGGGATCCGCTGGCTCCGCTGCTGGTTCTGGCGAA**TGA**GCATCCAAGGGC |
| #4 | pCTCON2-GFP-Rev | GAGCTCATCCATGCCATGAGTAATGCCTGC |
| #5 | Fwd_pRS315_NdeI_backbone | AACTCCTCAATCTGGTCGTTGGCTAACATATGTCACGC |
| #6 | Rev_LeuOmeRS-tRNA_TGA_Anticodon | CGGATGGTGGAATCGGTAGACACAAGGGATT**TCA**AATCCCTCGGCGTTCGCGCTGTGC |
| #7 | Fwd_LeuOmeRS-tRNA_TGA_Anticodon | GCACAGCGCGAACGCCGAGGGATT**TGA**AATCCCTTGTGTCTACCGATTCCACCATCCG |
| #8 | Rev_pRS315_XmaI_backbone | CCAGCTGGCAATTCCGGCTGTCAGCCCGGG |
| #9 | pCTCON2_Fwd_AltTGA-BXG | AAGCTTAATATGGGA**TGA**GCTGGCTCCGCTGCTGGTTCTGGCGAATACGCATCCAAGGGC |
| #10 | pCTCON2_Rev_AltTGA-BXG | GCCCTTGGATGCGTATTCGCCAGAACCAGCAGCGGAGCCAGC**TCA**TCCCATATTAAGCTT |
| #11 | Fwd_pRS315_MaPylRS_NdeI | GGAGCGAGTATCTCCTACCTAAATGGAGCGAAAATTAACTAACA |
| #12 | FwdpRS315_MaPylRS_TGA_Anticodon | GGTGTCGAACCCCGCTATGCTAGGTTT**TGA**AGACCCGCTGGTCGCCGGACCGTCC |
| #13 | RevpRS315_MaPylRS_TGA_Anticodon | GGACGGTCCGGCGACCAGCGGGTCT**TCA**AAACCTAGCATAGCGGGGTTCGACACC |
| #14 | Fwd_pRS315_OmeRS-7_NdeI | GCTGGAAATAACTCGAGTCATGTAATTACATATGTCACGC |
| #15 | RevpRS315_OmeRS-7_TGA | CCGAGCGGCCAAAGGGAGCAGACT**TCA**AATCTGCCGTCATCGACTTCGAAGGTTC |
| #16 | FwdpRS315_OmeRS-7_TGA | GAACCTTCGAAGTCGATGACGGCAGATT**TGA**AGTCTGCTCCCTTTGGCCGCTCGG |
| #18 | pRS416-BXG-EcoRI-Fwd | CTTTAACGTCAAGGAGAAAAAACCCCGGATCGAATTCC |
| #19 | pRS416-BglII-Rev | GTTACATCTACACTGTTGTTATCAGATCTCGAGC |
| #20 | pRS416-XmaI-Fwd | CAAAGACTGCAGGAATTCGATATCAAGCTTCC |
| #21 | pRS416-Linker-TGA-1st-Fwd | CACAAGCTTAATATGGGA**TGA**GCTGGCTCCGCTGCTGGTTCTGGCGAA**TAG**GCATCC |
| #22 | pRS416-Linker-TGA-1st-Rev | GGATGC**CTA**TTCGCCAGAACCAGCAGCGGAGCCAGC**TCA**TCCCATATTAAGCTTGTG |
| #23 | pRS416-Linker-TGA-2nd-Fwd | CAAGCTTAATATGGGA**TAG**GCTGGCTCCGCTGCTGGTTCTGGCGAA**TGA**GCATCCAAG |
| #24 | pRS416-Linker-TGA-2nd-Rev | CTTGGATGC**TCA**TTCGCCAGAACCAGCAGCGGAGCCAGC**CTA**TCCCATATTAAGCTTG |
| #25 | pCTCON2-PstI-Fwd | CCCATACGACGTTCCAGACTACGCTCTGCA |
| #26 | pCTCON2-XhoI_Rev | CGATTTTGTTACATCTACACTGTTGTTATCAGATCTCGAGC |
| #27 | Donkey1.1_L93TGA_Fwd | CTGCCAACAATCCTAC**TGA**ACTCCACCTACATTTGGTG |
| #28 | Donkey 1.1_L93TGA_Rev | CACCAAATGTAGGTGGAGT**TCA**GTAGGATTGTTGGCAG |

| | | |
|---|---|---|
| **#29** | pCTCON2-H54TGA-Fwd | GTTTCTGCGATATCTGGA**TGA**GGTGGGTCAACTTAC |
| **#30** | pCTCON2-H54TGA-Rev | GTAAGTTGACCCACC**TCA**TCCAGATATCGCAGAAAC |
| **#31** | pCTCON2-EcoRI-Fwd | CTTTAACGTCAAGGAGAAAAAACCCCGGATCG |
| **#32** | pCTCON2-SPS-Cmyc-GPD-Rev | GATAAACTGAGCTCCTCGAGCTATTACAAGTCCTCTTCAGAAATAAGCTTTTGTTC |
| **#33** | pCTCON2-SPS-Cmyc-GPD-Fwd | GAACAAAAGCTTATTTCTGAAGAGGACTTGTAATAGCTCGAGGAGCTCAGTTTATC |
| **#34** | pCTCON2-SPS-NcoI-Rev | GCTCTTGCAATTGTTTAATCAAGTTACTGCTTGCCATGG |
| **#35** | pCHA-OTS-AcFRS-New-Fwd | CATAAACACACAGTATGTTTTTAACTCGAGGAGCTCAGTTTATC |
| **#36** | pCTCON2-TyrAcFRS-NdeI-Rev | GAGGGCGTGAATGTAAGCGTGACATATGTAATTACATGAC |
| **#37** | pCTCON2-TyrAcFRS-NdeI-Fwd | TTACATATGTCACGCTTACATTCACGCCCTC |
| **#38** | pCHA-OTS-TyrAcFRS-new-Rev | TTGTAATACGACTCACTATAGGGCGAATTGGAGCTC |
| **#39** | pCHA-SPS-EcoRI-Fwd | GATCGAGGTCGACGGTATCGATGA |
| **#40** | pCHA-SPS-XmaI-Rev | GTGGACAAGTATGGGTTTTGTCGG |

*Mutations incorporated within the primer sequences for changing the orthogonal codon or anticodon sequence or for incorporating an additional orthogonal codon are highlighted with different colors.

**Table S2.** Vector backbones for all the reporters and OTSs used in this study and their selection markers in yeast and *E. coli*.

| Reporter/OTS Name | Vector Backbone | Selection Marker for Yeast | Antibiotic Resistance Marker for *E. coli* |
|---|---|---|---|
| BYG (For RJY100) | pCTCON2 | Trp1 | Amp |
| BXG-TAG or BXG-TGA (For RJY100) | | | |
| Alt-TAG or Alt-TGA (For RJY100) | | | |
| Donkey1.1-WT | | | |
| Donkey1.1-AO-TyrAcFRS/$tRNA_{CUA}^{Tyr}$ | | | |
| Donkey1.1-OA-TyrAcFRS/$tRNA_{CUA}^{Tyr}$ | | | |
| T2RS5/$tRNA^{Tyr}$ | pRS315 | Leu2 | Kan |
| LeuOmeRS/$tRNA^{Leu}$ | | | |
| MaPylRS/$tRNA^{Pyl}$ | | | |
| LysAlkRS3/$tRNA_{UCA}^{leu}$ | | | |
| BYG (For BY4741) | pRS416 | Ura3 | Amp |
| BXG-TAG or BXG-TGA (For BY4741) | | | |
| Alt-TAG or Alt-TGA (For BY4741) | | | |
| BYG-TyrOmeRS/$tRNA_{CUA}^{Tyr}$ | | | |
| $BX_2G$-AO-TyrOmeRS/$tRNA_{CUA}^{Tyr}$ | | | |
| $BX_2G$-OA-TyrOmeRS/$tRNA_{CUA}^{Tyr}$ | | | |
| Donkey1.1-OA-TyrAcFRS/$tRNA_{CUA}^{Tyr}$ (For protein purification) | pCHA | Trp1 | Amp |

*Trp- Tryptophan; Amp- Ampicillin; Kan- Kanamycin; Leu - Leucine

**Table S3.** List of yeast strains and plasmid combinations used in this study along with the selection media employed for their propagation and induction.

| Yeast Strains | Plasmid Combinations Used | Selection Media for Propagation and Induction |
|---|---|---|
| RJY100 | i) pCTCON2-BYG, or<br>ii) pCTCON2-BXG-TAG, or<br>iii) pCTCON2-BXG-TGA, or<br>iv) pCTCON2-AltTAG, or<br>v) pCTCON2-AltTGA, or<br>+<br>i)pRS315-LeuOmeRS/tRNA*$^{Leu}$, or<br>ii) pRS315- MaPylRS/tRNA*$^{Pyl}$, or<br>iii) pRS315-T2RS5/tRNA*$^{Tyr}$<br>(*CUA or UCA tRNA anticodon depending on the experimental conditions) | SD-SCAA -Trp -Leu -Ura;<br>SG-SCAA -Trp -Leu -Ura |
| WT BY4741<br>*ppq1Δ* BY4741<br>*tpa1Δ* BY4741 | i) pCTCON2-BYG, or<br>ii) pCTCON2-BXG-TAG, or<br>iii) pCTCON2-BXG-TGA, or<br>iv) pCTCON2-AltTAG, or<br>v) pCTCON2-AltTGA, or<br>+<br>i)pRS315-LeuOmeRS/tRNA*$^{Leu}$, or<br>ii) pRS315- MaPylRS/tRNA*$^{Pyl}$, or<br>iii) pRS315-T2RS5/tRNA*$^{Tyr}$<br>(*CUA or UCA tRNA anticodon depending on the experimental conditions) | SD-SCAA -Leu -Ura;<br>SG-SCAA -Leu -Ura |
| WT BY4741<br>*ppq1Δ* BY4741 | i)pRS416-BYG-TyrOmeRS /$tRNA_{CUA}^{Tyr}$, or<br>ii) pRS416-B$X_2$G-AO-TyrOmeRS /$tRNA_{CUA}^{Tyr}$, or<br>iii) pRS416-B$X_2$G-AO-TyrOmeRS /$tRNA_{CUA}^{Tyr}$<br>+<br>i) pRS315-LeuOmeRS/$tRNA_{UCA}^{Leu}$, or<br>ii) pRS315- MaPylRS/$tRNA_{UCA}^{Pyl}$ | SD-SCAA -Leu -Ura;<br>SG-SCAA -Leu -Ura |
| RJY100 | i) pCTCON2-Donkey1.1-WT, or<br>ii) pCTCON2- Donkey1.1-AO-TyrAcFRS/$tRNA_{CUA}^{Tyr}$, or<br>iii) pCTCON2- Donkey1.1-AO-TyrAcFRS/$tRNA_{CUA}^{Tyr}$, or<br>+<br>pRS315-LysAlkRS3/$tRNA_{UCA}^{Leu}$ | SD-SCAA -Trp -Leu -Ura;<br>SG-SCAA -Trp -Leu -Ura |

**Table S4.** DNA sequences of 'Linker' region in dual fluorescent reporter construct and scFv reporters with orthogonal codon incorporation sites highlighted.

| Name | DNA Sequence (5'→3')* |
|---|---|
| Linker Wild-type (BFP-GFP construct) | ATGGGATCCGCTGGCTCCGCTGCTGGTTCTGGCGAATAC |
| Donkey1.1-WT | CTGCAGGCTAGTGGTGGAGGAGGCTCTGGTGGAGGCGGTAGC<br>GGAGGCGGAGGGTCGGCTAGCGACATACAGATGACTCAAAGT<br>CCCAGTTCACTATCTGCGTCTGTTGGTGATAGAGTCACCATTAC<br>GTGTAGAGCTTCTCAGTCGATTAGCTCGTACTTGAATTGGTATC<br>AACAGAAACCAGGGAAAGCTCCAAAGTTGCTGATCTATGCAGC<br>ATCTAGCTTACAAAGTGGTGTACCTTCCAGGTTTTCAGGCTCAG<br>GATCTGGAACTGATTTCACACTTACCATATCATCCTTACAACCG<br>GAAGATTTCGCCACATATTACTGCCAACAATCCTACTCTACTCC<br>ACCTACATTTGGTGGTGGCACTAAAGTGGAGATTAAGGGTACT<br>ACTGCCGCTAGTGGTAGTAGTGGTGGCAGTAGCAGTGGTGCCG<br>AGGTGCAATTGCTAGAATCAGGAGGTGGTTTGGTACAACCTGG<br>TGGTAGCTTAAGGTTGTCTTGTGCTGCTAGTGGATTCACGTTTA<br>GTAGCTATGCCATGTCATGGGTTAGACAAGCTCCAGGTAAAGG<br>CTTAGAATGGGTTTCTGCGATATCTGGATCTGGTGGGTCAACTT<br>ACTATGCAGATTCCGTCAAAGGCAGATTTACCATTTCCAGAGA<br>CAATTCGAAGAATACACTGTACCTTCAGATGAACTCGTTACGT<br>GCAGAAGATACTGCTGTTTACTACTGTGCTAAGTATGACAAGA<br>CGCACCACAATCCTGATTACGCTCTCGACTACTGGGGCCAAGG<br>AACCCTGGTCACCGTCTCCTCAGGATC |

* The highlighted region indicates the sites where TAG or TGA codons were introduced into DNA encoding the reporters.
